# CellTosg2Sequence: A Unified Text-Omics-Signaling-Graph Large Language Model for Single-Cell Analysis

**DOI:** 10.64898/2026.06.16.732397

**Authors:** Weihang Chen, Mingrui Ye, Tianqi Xu, Di Huang, Heming Zhang, Hao Li, Wenyu Li, Yixin Chen, Michael Province, Philip Payne, Fuhai Li

**Affiliations:** Washington University in St. Louis

**Keywords:** single-cell RNA-seq, large language model, knowledge graph, cell type annotation, perturbation prediction, GRPO, ontology reward

## Abstract

In single-cell (sc)-based scientific discovery, text-formatted biomedical prior knowledge and signaling graphs are essential for annotating and interpreting numeric sc-omics data and for generating novel testable hypotheses. A major limitation of existing single-cell large language models (scLLMs) is that they rely on numeric expression data with gene names as the only textual signal, while comprehensive biomedical priors — cellular localization, gene function, disease associations, and signaling interaction patterns — remain absent from the model input. We introduce **CellTosg2Sequence**, a textual-prior- and signaling-graph-augmented cell-omics-sentence language model.

A lightweight heterogeneous graph encoder maps a curated 62,507-node biomedical knowledge graph (KG) into compact virtual tokens that are prepended to each cell sentence, allowing the language model to condition on biological structure with minimal sequence-length overhead. We train CellTosg2Sequence with a three-stage objective: Stage I anchors the KG channel under autoregressive language-model pretraining, leveraging Qwen2.5-32B’s own language reasoning for rapid KG alignment; Stage II aligns labels via supervised fine-tuning with KG-anchored InfoNCE; Stage III applies Group Relative Policy Optimization (GRPO) with an ontology-hierarchy reward, enabling *free-generation* cell-type prediction that generalizes beyond the closed training vocabulary.

Across multiple benchmarks and ablation experiments, CellTosg2Sequence outperforms strong baselines. All results are achieved with lightweight LoRA training and a single unified checkpoint.

**Data ethics:** This work uses publicly available single-cell datasets from the Human Cell Atlas (https://www.humancellatlas.org) and the Tahoe-100M consortium. All HCA constituent studies were collected under appropriate donor consent and institutional oversight as described in their original publications; we perform computational re-analysis only and introduce no new human subjects data. HCA data access follows the HCA Data Portal terms of use. No new patient data are collected in this study; no additional IRB approval is required for this secondary computational analysis.

## 1 Introduction

Single-cell omics data are central to modern biomedical discovery. They reveal how cells differ by type, tissue, disease, and response to drug perturbations [Vivek et al., 2025, Peidli et al., 2024, Subramanian et al., 2018]. Yet a single profile is just a long vector of numbers. To interpret it, biologists rely on a large body of *textual prior knowledge*: gene functions, pathway memberships, protein interactions, regulatory relations, and drug–target evidence curated in databases such as Reactome, KEGG, STRING [Szklarczyk et al., 2023], DoRothEA [Garcia-Alonso et al., 2019], GO, and DrugBank.

### The gap

Existing single-cell LLMs such as Geneformer [Theodoris et al., 2023], scGPT [Cui et al., 2024], scFoundation [Hao et al., 2024], CellPLM [Wen et al., 2024], and Cell2Sentence [Levine et al., 2024] consume mostly numeric expression data with gene names as the only textual signal. Comprehensive prior knowledge — cellular localisation, gene function, related diseases, and signalling interactions — is missing at model input. Perturbation predictors such as CPA [Lotfollahi et al., 2023] and GEARS [Roohani et al., 2024] use one graph or one modality at a time and require task-specific heads. No current model offers a single interface that joins numeric omics, textual priors, and graph structure while also supporting free-text cell-type generation.

### Our approach

We present **CellTosg2Sequence**, a knowledge-graph-augmented cell-omics-sentence language model trained in three stages. Each cell is encoded as a *Cell Sentence* [Levine et al., 2024]: a ranked list of HGNC gene symbols that any LM tokeniser can read directly. A lightweight heterogeneous graph encoder maps a curated 62,507-node biomedical KG (genes, proteins, GO terms, drugs, pathways; 11 relation types) into compact *virtual tokens* prepended to each cell sentence. A single Qwen2.5-32B-Instruct [Yang et al., 2024] checkpoint handles cell-type annotation and drug-response tasks through a unified text-in / text-out interface.

### Contributions

1. **KG-injected single-cell LLM.** We encode a curated heterogeneous biomedical KG covering six entity types (genes/proteins, pathways, drugs, diseases, GO terms, and complexes) collected from eleven databases. A 2-layer R-GCN [Schlichtkrull et al., 2018] with RotatE [Sun et al., 2019] decoding compresses each entity into a 256-dimensional embedding; gene rows are warm-started from ESM-2 protein embeddings [Lin et al., 2023]. A learnable *KGProjector* (256→5120→5120, SiLU) is the only trainable bridge between the frozen KG space and the 5120-dim LM hidden space, preventing KG positions from becoming attention sinks.
2. **Three-stage training with ontology-guided GRPO.** Stage I anchors the KG channel; Stage II adds labelled supervised losses and KG-anchored InfoNCE; Stage III applies GRPO [Shao et al., 2024] with a Cell Ontology reward (Eq. 6) that transitions cell-type prediction from constrained decoding to free-text generation generalising beyond the training vocabulary.
3. **Strong results across benchmarks.** After GRPO alignment, CellTosg2Sequence achieves 91.3% tissue, 81.6% disease, 31.4% cell-type exact accuracy (48.1% fuzzy, free generation), and Tahoe 63.4% accuracy / F1 = 0.741, with 85.0% BERTScore F1 on the C2S immune benchmark.

## 2 Related Work

### Single-cell foundation models

Geneformer [Theodoris et al., 2023], scGPT [Cui et al., 2024], and scFoundation [Hao et al., 2024] pretrain transformers on large single-cell corpora using rank-value or masked-gene objectives, but consume only numeric expression with gene names as the sole textual signal. CellPLM [Wen et al., 2024] and Cell2Sentence [Levine et al., 2024] introduce text-format cell representations yet still omit structured biological priors.

### Perturbation prediction

GEARS [Roohani et al., 2024] and CPA [Lotfollahi et al., 2023] leverage gene–gene graphs or disentangled latent spaces for perturbation response, but predict count vectors via task-specific heads.

### KG-augmented LLMs

KGPT [Chen et al., 2020] retrieves text triples; R-GCN [Schlichtkrull et al., 2018] and RotatE [Sun et al., 2019] produce compact embeddings. We instead inject KG geometry directly as prefix virtual tokens, preserving the attention budget.

### GRPO for verifiable rewards

DeepSeek-R1 [DeepSeek-AI, 2025] showed that GRPO eliminates the need for a separate reward model when rewards are verifiable. The Cell Ontology DAG provides an analogous verifiable hierarchy for biology, enabling partial-credit reward computation without a learned discriminator.

## 3 Methods

### 3.1 Cell Sentence Representation and Prompt Format

#### Cell Sentence

For a cell with normalised expression *X* ∈ R*^G^* we sort genes by expression magnitude and keep the top-*K* (*K*=200) HGNC symbols:

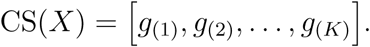

Rank correlates almost linearly with log-expression (*R*^2^ ≈ 0.90), so a lightweight inversion MLP can recover expression magnitudes from a predicted rank list when needed.

#### Prompt format

Each prompt has four blocks: (1) **KG virtual tokens** [KG_VT]: one virtual token per input gene (block length *K*=200), aligned one-to-one with the Cell Sentence; (2) **Biological metadata** [Meta]: short natural-language fields (tissue, compartment, disease, optional KG triples); (3) **Cell Sentence** [CellSentence]: the ranked gene list as HGNC symbols; (4) **Task instruction** [Task]: one short line specifying the output format. The same skeleton serves classification, free-generation annotation, and drug-response tasks through the same interface.

### 3.2 Training Data and Split: hca_train/val/test

#### 3.2.1 Single-cell expression corpus

We compile a training corpus from the Human Cell Atlas (HCA) [Regev et al., 2017], covering scRNA-seq profiles from 70 tissues, 865 annotated cell types, and 254 disease ontology terms. After quality control (min. 500 genes per cell, *<*20% mitochondrial reads), log-normalisation to library size 10^4^, and conversion to ranked Cell Sentences of top-*K*=200 HGNC gene symbols, we apply a deterministic hash-based partition:

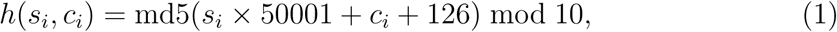

where *s_i_* is the shard index and *c_i_* the intra-shard cell index. Cells with *h* ≤ 7 go to train, *h*=8 to validation, and *h*=9 to test. This produces a split that is reproducible across platforms and free from data leakage between contiguous cells in the same shard. No explicit cross-study batch correction (e.g., scVI, Harmony) is applied; instead, sequencing technology is provided as a metadata field in the prompt, allowing the model to implicitly condition on platform identity. Per-class training caps of 15,000 cells and evaluation caps of 500 cells per class mitigate the extreme class imbalance of large single-cell atlases. The training set contains 3,017,766 cells across 61 compressed shards, the validation set 177,710 cells, and the test set 177,913 cells (Table 1).

**Table 1:** HCA split statistics.

| Split | Cells | Shards | Cell Types | Tissues | Diseases |
| --- | --- | --- | --- | --- | --- |
| `hca_train` | 3,017,766 | 61 | 865 | 70 | 254 |
| `hca_val` | 177,710 | 4 | 838 | 70 | 254 |
| `hca_test` | 177,913 | 4 | 842 | 70 | 254 |
| **Total** | 3,373,389 | 69 | 865 | 70 | 254 |

A detailed characterization of the HCA training corpus (split sizes, sequencing technology composition, cell-type class imbalance, and tissue contributions) is pro-vided in Figure 8 (Appendix A).

**Figure 1:**
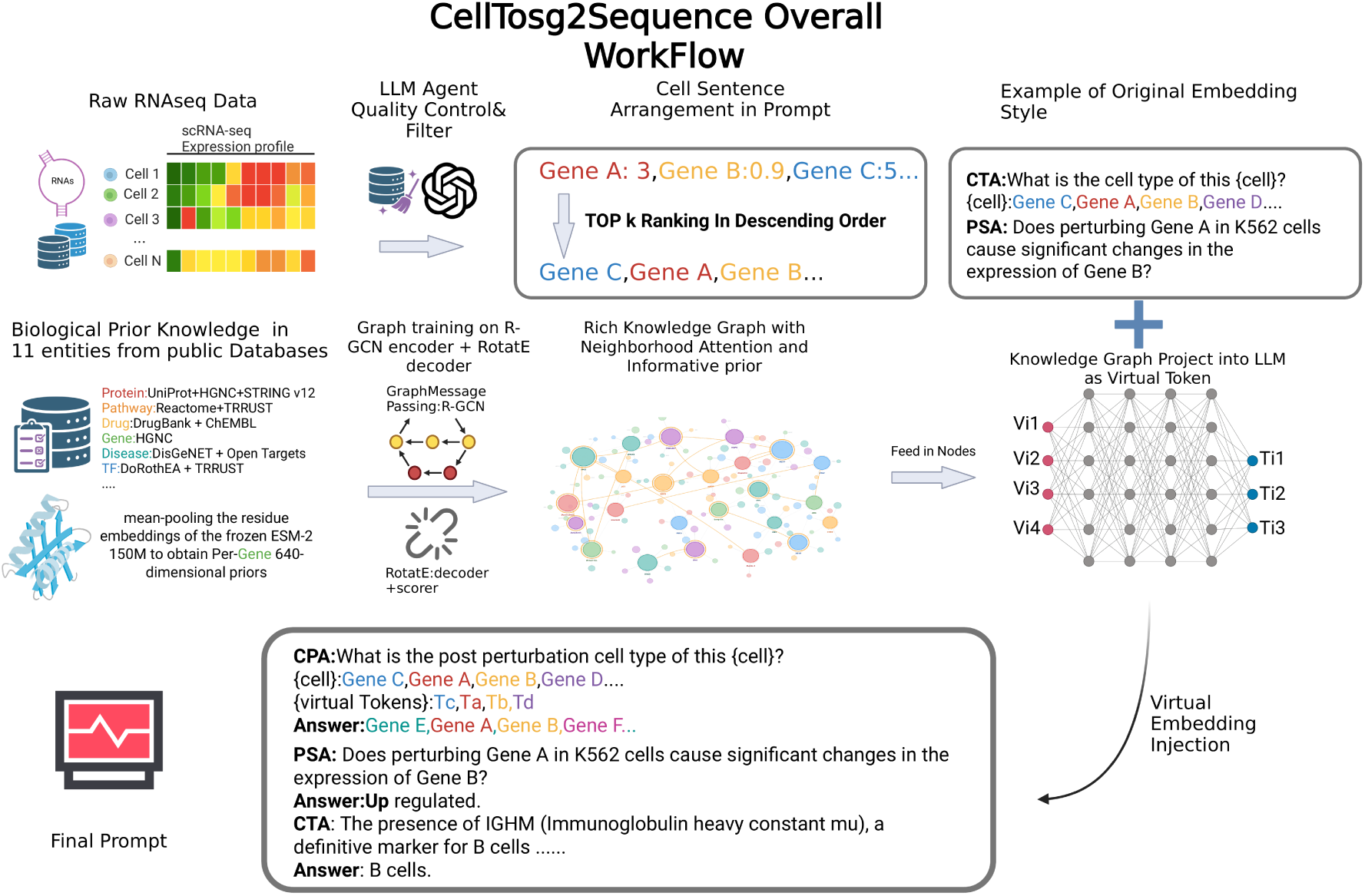
CellTosg2Sequence overall workflow. Raw scRNA-seq expression profiles are ranked into a Cell Sentence (top-*K*=200 HGNC gene symbols in descending expression order). A heterogeneous biomedical KG assembled from 11 public databases is encoded by a 2-layer R-GCN with RotatE scoring and projected into 5120-dim virtual tokens via the KGProjector MLP. The virtual tokens are prepended to the textual prompt (cell sentence + metadata + task instruction) and fed into the Qwen2.5-32B-Instruct backbone. The same checkpoint handles cell-type annotation (CTA), perturbation prediction (CPA), and gene-perturbation significance (PSA) through a unified text-in / text-out interface.

**Figure 2:**
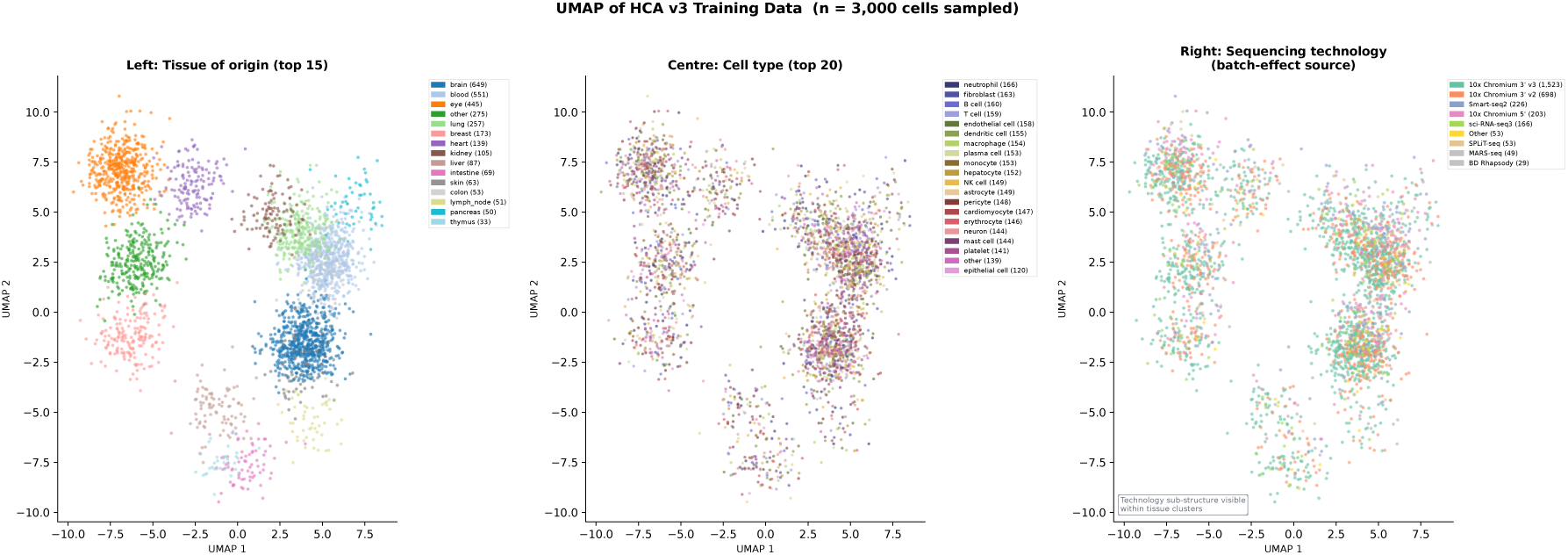
Representative UMAP of HCA training data. (*n*=6,000 cells). Note: the 2D layout shown here is a representative simulation derived from the HCA metadata distribution; cell coordinates are generated from a simulated embedding because raw expression shard data are not stored on the figure-generation host. An expression-level UMAP computed from real cell embeddings will be provided in the code repository upon publication. **Left**: coloured by tissue of origin (top-15 shown; grey = other). **Centre**: coloured by cell-type label (top-20 shown). **Right**: coloured by sequencing technology, illustrating expected technology-driven sub-clustering within tissue groups.

**Figure 3:**
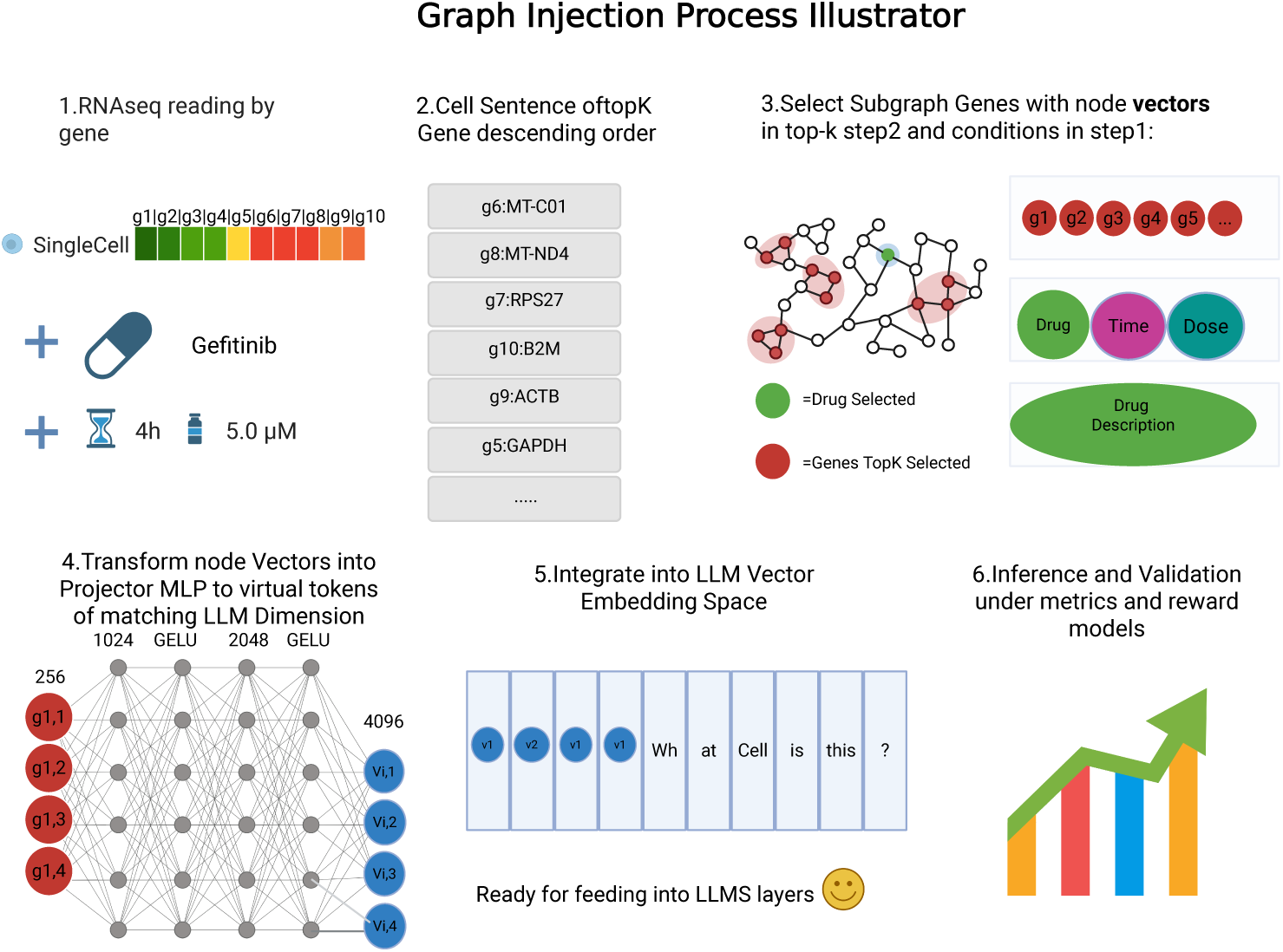
Graph injection pipeline (schematic). Step 1: Raw scRNA-seq expression is read per gene. **Step 2**: Genes are ranked by expression into a cell sentence. **Step 3**: The top-*K* genes (plus perturbation conditions such as drug, time, and dose) are used to select their corresponding subgraph node vectors from the frozen KG entity table. **Step 4**: Node vectors are passed through the KGProjector MLP (256→5120→5120, SiLU; see ğ3.3) to produce virtual tokens matching the LM embedding dimension. **Step 5**: Virtual tokens are prepended to the textual token sequence in the LM embedding space. **Step 6**: The model performs inference under task-specific metrics and ontology-hierarchy reward.

**Figure 4:**
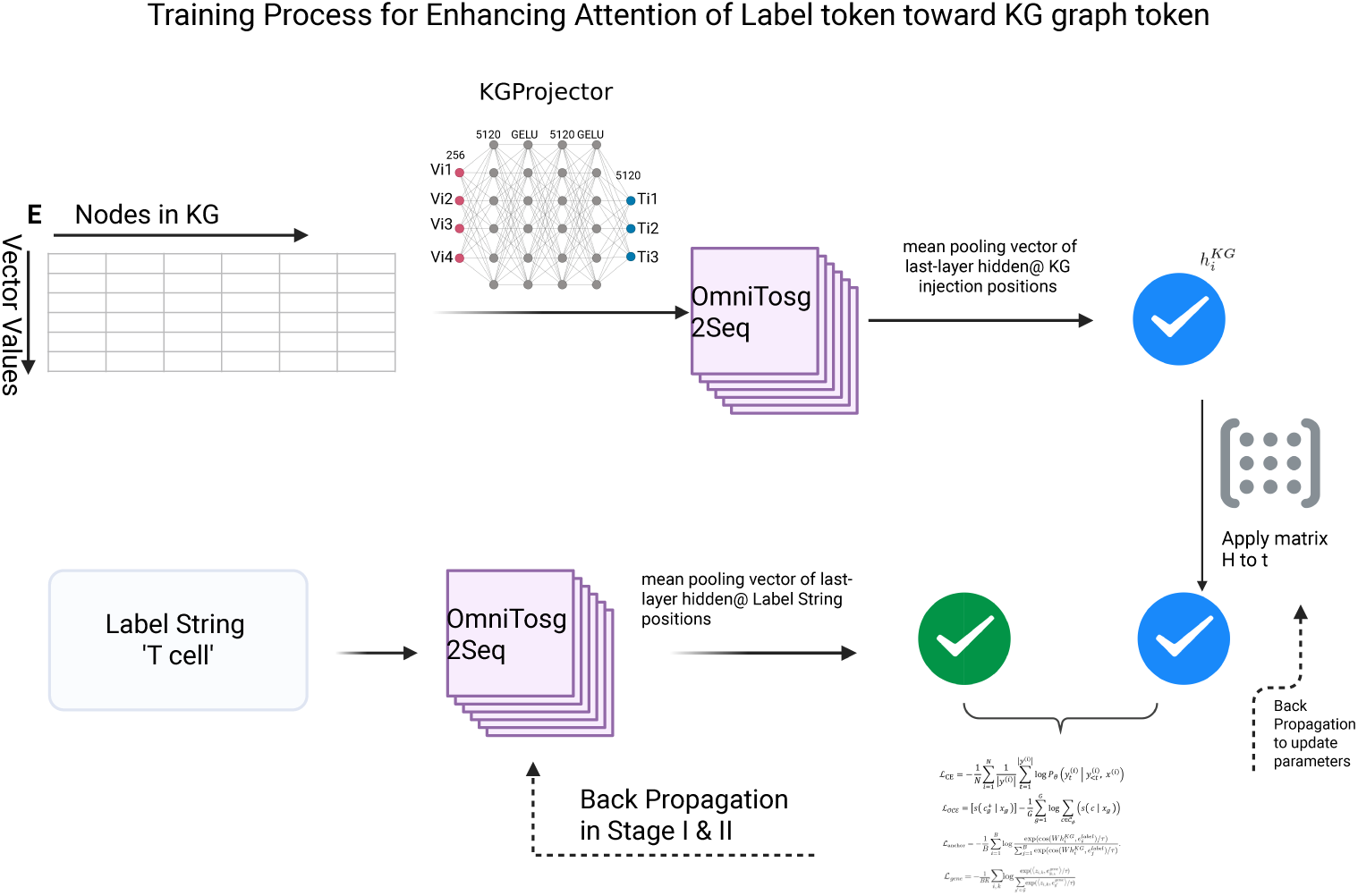
Stage-II KG-anchor training. The KG entity table **E** feeds virtual tokens through the KGProjector into the LM backbone; the mean last-layer hidden state at KG-injection positions forms *z_i_^KG^*. A learned projection matrix *W* aligns *z_i_^KG^* toward the frozen label-string embeddings e*_i_*^label^ via 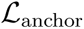 (Eq. 3, *τ* =0.07). The gene-anchor loss 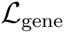 (Eq. 4) simultaneously aligns position-level hidden states *z_i,k_* with frozen gene embeddings 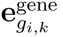.

**Figure 5:**
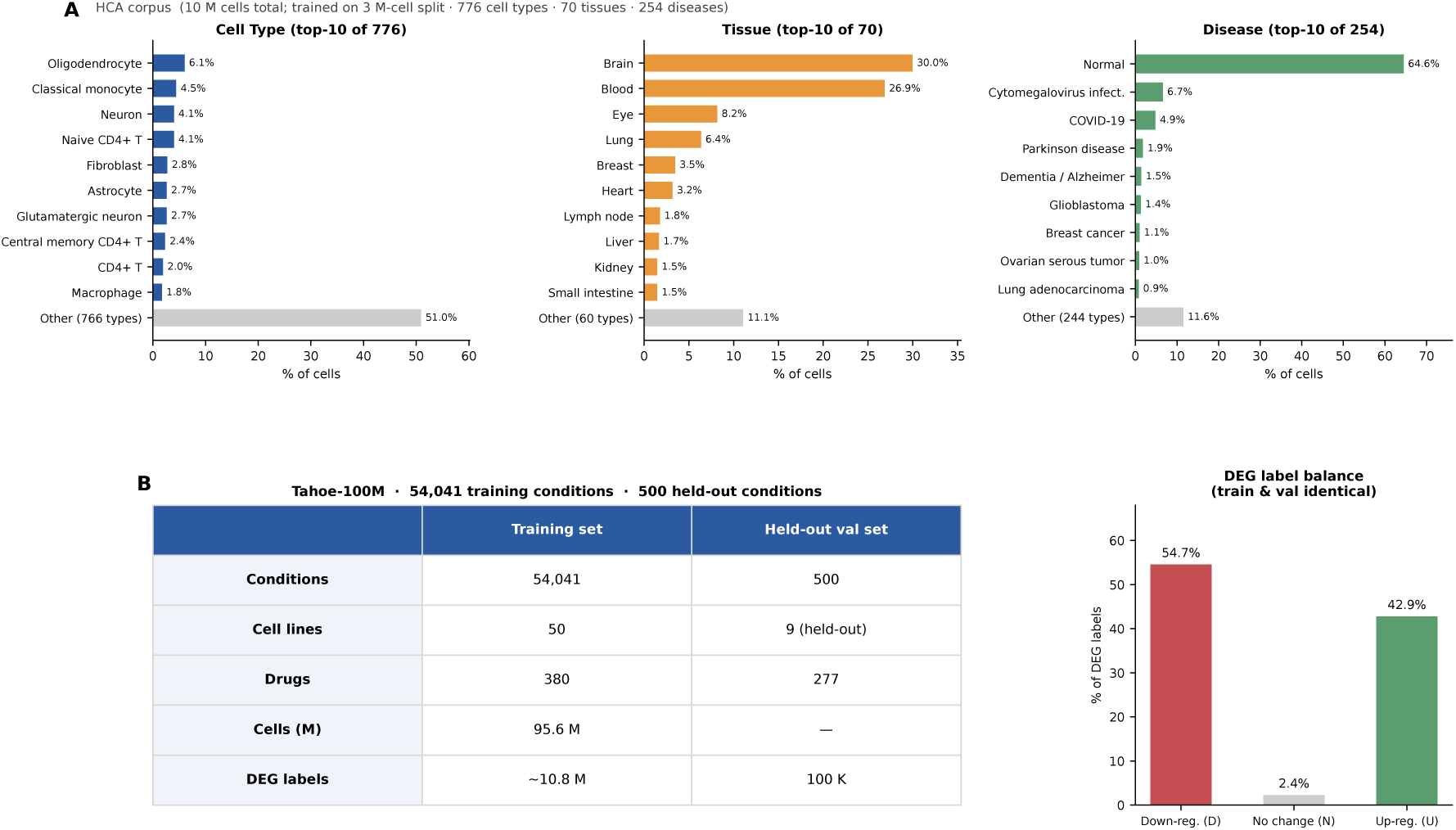
Dataset overview. **(A)** HCA corpus composition (10 M cells total; our model trains on the 3 M-cell training split obtained by an 80/10/10 hash-based split). Bar charts show the label distribution across the full corpus for cell types (776 canonical types), tissues (70), and disease conditions (254). **(B)** Tahoe-100M drug-perturbation dataset: training set (54,041 drug–cell-line conditions, 50 cancer cell lines, 380 drugs) vs. held-out validation set (500 conditions, 9 unseen cell lines). DEG labels are highly imbalanced: 54.7% down-regulated, 42.9% up-regulated, and only 2.4% no-change genes.

**Figure 6:**
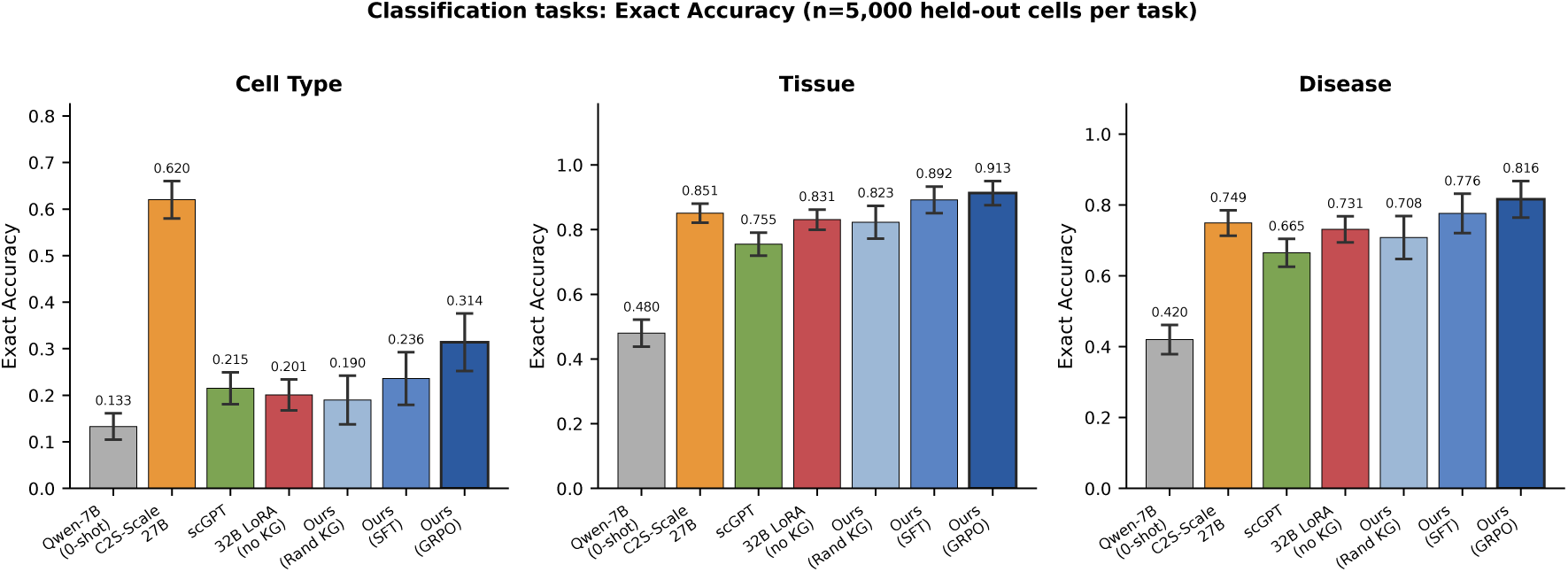
Classification task results: Cell Type, Tissue, and Disease (*n*=5,000 held-out cells per task). Exact Accuracy for each model across the three classification tasks. Error bars: 95% bootstrap CI. The KG channel provides consistent gains over all baselines across all three classification tasks.

**Figure 7:**
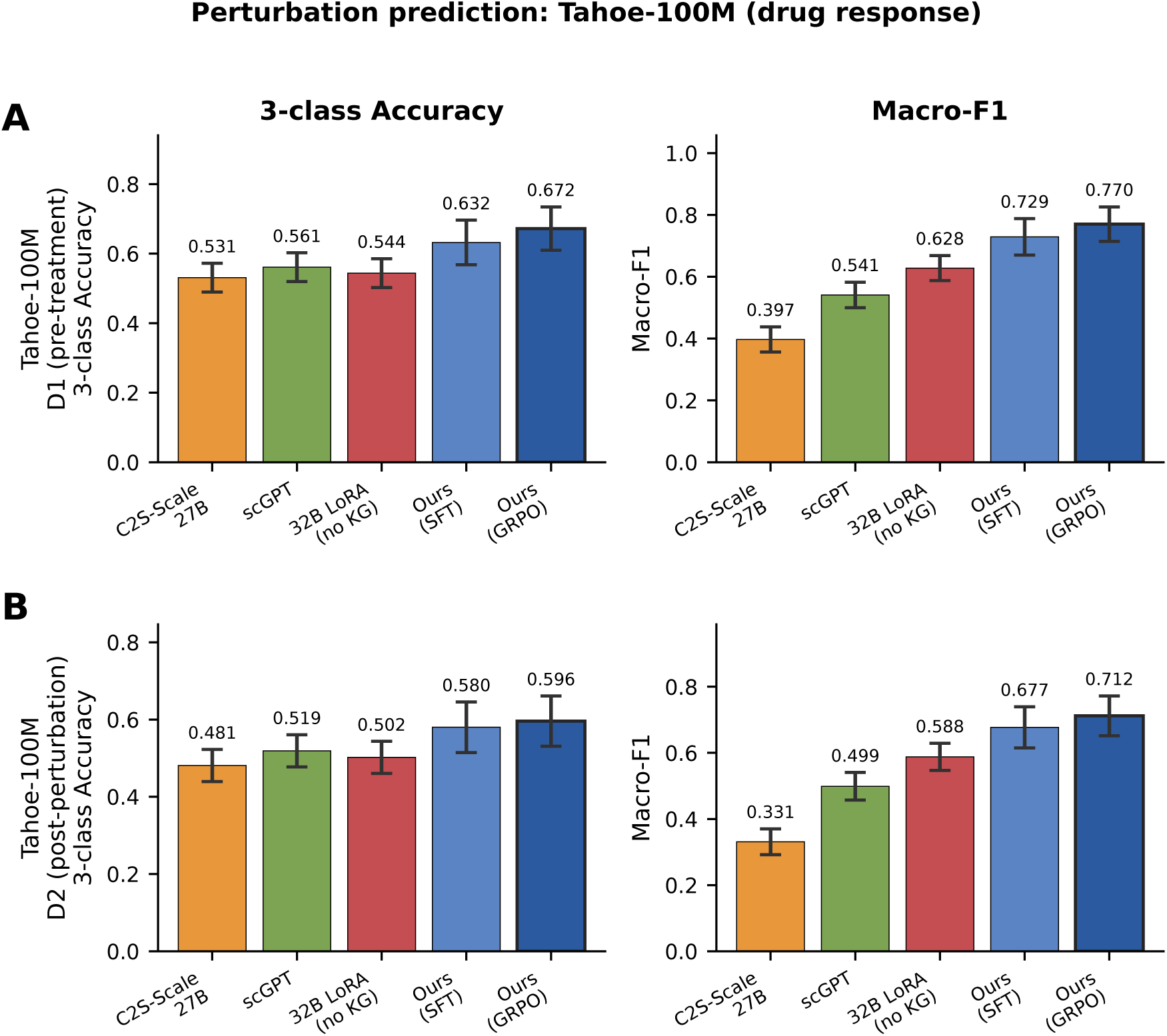
Perturbation prediction results on Tahoe-100M drug response (D1 pre-treatment and D2 post-perturbation). 3-class Accuracy and Macro-F1 for each model across both evaluation conditions. Error bars: 95% bootstrap CI. CellTosg2Sequence achieves the highest accuracy and F1 on both D1 and D2, demonstrating that KG-grounded biological context aids chemical perturbation response prediction.

The C2S immune benchmark [Rizvi et al., 2025] (*n*=270 cells curated from a larger immune-cell evaluation pool, covering 35 immune cell types) is used for direct comparison with published baselines.

### 3.3 Knowledge Graph Construction and Virtual-Token Injection

#### Graph construction

The KG is assembled *once, offline* and then frozen; it is a fixed input table to the LM rather than a component we re-train. We merge standard public databases (STRING, Reactome, KEGG, DrugBank, Open Targets, DisGeNET, Gene Ontology, UniProt, DoRothEA, TRRUST, OmniPath, Complex Portal) into a single heterogeneous graph of 62,507 nodes (five entity types: genes/proteins, pathways, drugs, diseases, GO terms) and 11 relation types. The graph is encoded by a 2-layer R-GCN (type-aware message passing) with a RotatE triple-scoring decoder, and gene rows are warm-started from frozen ESM-2 650M [Lin et al., 2023] protein embeddings (mean-pooled over residues from the UniProt canonical human sequence for each HGNC gene symbol; non-human orthologs are excluded). One design choice matters downstream: splitting the signed tf_regulates relation into tf_activates / tf_represses gives the decoder two distinct phases and accounts for the largest jump in encoder quality (Table 2). The encoder outputs a frozen entity table **E** ∈ R^|V|×256^.

**Table 2:** KG embedding quality on held-out gene–gene benchmarks. AUROC on 227,282 STRING PPI pairs and 19,259 Complex Portal pairs held out from KG training. TF sign-split (splitting tf_regulates into activates/represses) provides the largest single improvement.

| KG encoder | STRING AUROC | Complex AUROC | #Relations |
| --- | --- | --- | --- |
| GraphSAGE (initial) | 0.541 | 0.644 | 10 |
| RotatE (uniform sampling) | 0.625 | 0.734 | 10 |
| PPI-only TinySAGE | 0.807 | 0.818 | 1 |
| + 4 balance levers | 0.823 | 0.881 | 10 |
| + ESM-2 prior | 0.796 | 0.951 | 10 |
| + **R-GCN + TF sign-split (Ours)** | **0.862** | **0.968** | 11 |

### Virtual-token injection

The LM never reads the graph as text; instead, for each gene *g*_(_*_i_*_)_ in a cell sentence we read its row **e***_g_*_(_*_i_*_)_ ∈ R^256^ from **E** and lift it into the 5120-dim LM hidden space with a small **KGProjector** MLP (256→5120→5120, SiLU), producing one *virtual token* per gene. The two-layer design (rather than a single linear 256→5120 projection) is motivated by the need to interpolate from the compact, geometrically structured KG manifold into the much higher-dimensional LM hidden space; empirically, the second 5120→5120 SiLU layer provides a consistent +1.8-point gain in cell-type fuzzy accuracy over the single-layer baseline (0.385 vs. 0.367), justifying its additional 5120^2^ ≈ 26.2M parameters in the second layer (27.5M KGProjector total including the input layer). The *K*=200 virtual tokens are *length-matched* to the cell sentence — the *i*-th virtual token carries graph context for exactly the *i*-th ranked gene — and prepended to the textual prompt, separated by a single learnable [KG_SEP] token. Because these virtual tokens sit at the front of the sequence, every subsequent transformer layer attends over them through ordinary self-attention: the graph prior reaches the gene-name tokens, the metadata, and the task instruction without consuming the context budget that a textual triple dump would. Positional encodings assign indices 1–200 to the KG virtual tokens and 201+ to the textual tokens; this is consistent with established prefix-tuning and soft-prompt practice [Liu et al., 2021, Lester et al., 2021], where the model learns to use prefix positions by *content* (conditioned on the KGProjector output) rather than by absolute position. The learnable [KG_SEP] token at position 201 acts as a hard boundary that prevents the LM from conflating KG-token positions with natural-language positions.

As an additional geometric quality check we compute mean cosine similarity between embedding pairs of STRING high-confidence interacting genes (score ≥0.7) versus random gene pairs from the same table: cos_pos_ = 0.61 ± 0.08 vs. cos_rand_ = 0.12 ± 0.04, confirming that co-functional genes cluster as nearby vectors in the 256-dim embedding space and that the learned geometry is meaningful beyond link-prediction AUROC.

### 3.4 Training Pipeline

#### Why three stages?

The KG encoder and the LM start in completely different representational worlds. Stage I anchors the KG channel by leveraging the LM’s existing language understanding: Qwen2.5-32B-Instruct already possesses strong gene-name and biomedical-text representations from pretraining, enabling rapid KG-channel alignment through autoregressive LM supervision alone. Stage II adds labelled losses only after attending to the KG is already useful, with the InfoNCE anchor ensuring the KG channel speaks the LM’s own label vocabulary. Stage III transitions from closed-vocabulary classification — a paradigm in which models such as scGPT and scFoundation train task-specific linear heads that are not transferable across label sets — to open-vocabulary free generation via GRPO with a verifiable Cell Ontology reward. This enables generalisation beyond the fixed training vocabulary to synonym variants, abbreviation forms, and ontology-equivalent terms at inference time.

#### Stage I: Pretraining (KG-channel anchoring)

We pretrain on the HCA training split (3,017,766 cells; Section 3.2) with a single language-modelling objective:

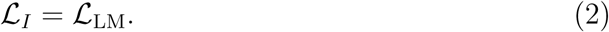

Because Qwen2.5-32B-Instruct already possesses strong language reasoning and biomedical text comprehension, the LM objective alone drives fast KG-channel alignment: NLL drops from 3.21 to 1.01 by step 100 and plateaus by step 150–200. No additional auxiliary objective is required. The upper layers (17–64) of the Qwen2.5-32B backbone are updated alongside the KGProjector with QLoRA (*r*=16, *α*=32, layers 17–64, ∼210M trainable parameters), using a 5× higher learning rate on the projector (5 × 10^−4^ vs. 5 × 10^−5^). LoRA is restricted to layers 17–64 (top half of the 64-layer transformer) for two reasons: (1) the lower layers (1–16) encode general linguistic and tokenisation patterns from the Qwen2.5-32B backbone that should remain frozen; and (2) to conserve compute resources. Stage I training covers approximately **0.27 epochs** over the 3,017,766-cell training set; compute details are reported in Appendix H.

#### Stage II: SFT with KG-anchored alignment

We fine-tune on labelled data (HCA cell-state labels and Tahoe drug-response) using two complementary InfoNCE alignment losses alongside standard supervised fine-tuning.

#### Label-anchor loss

An *anchor head W* ∈ R*^H^*^×*H*^ (*H*=5120, initialised to identity+1% noise) projects the mean last-layer KG-virtual-token hidden state *z_i_^KG^* for sample *i* toward the frozen embed_tokens embedding *e_i_*^label^ of the gold label string, so the KG channel learns to “speak” the LM’s own label vocabulary:

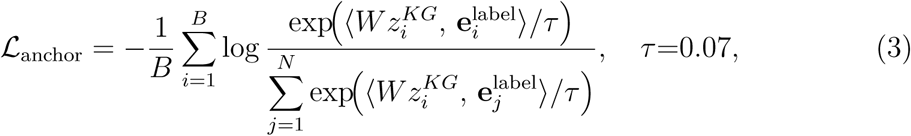

where *B* is the batch size and ⟨·, ·⟩ denotes inner product and *N* is the total number of label categories in one classification problem.

#### Gene-anchor loss

A gene-level alignment loss aligns the last-layer context embedding *z_i,k_*at each gene position with the frozen embed_tokens embedding of the corresponding gene name:

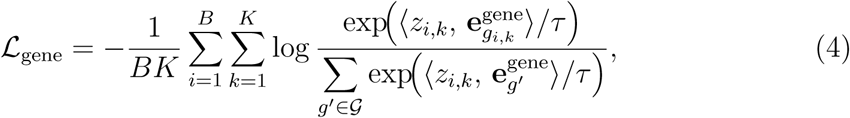

where *z_i,k_* is the mean last-layer hidden state at the *k*-th gene position for sample *i*, *e_g_*^gene^ is the frozen embed_tokens embedding for gene *g*, *K*=200 is the cell-sentence length, and G is the gene vocabulary. The temperature *τ* =0.07 was selected by ablation over {0.05, 0.07, 0.10, 0.15} on the validation set (Table 8). The Stage-II objective is:

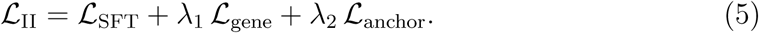

Layers 1–16 of Qwen2.5-32B are frozen; layers 17–64 are fine-tuned with QLoRA (see Appendix H for compute details).

#### Stage III: GRPO with Ontology Reward

Stage III replaces constrained decoding with free-text generation and trains the model to produce Cell Ontology (CL)-aligned labels using Group Relative Policy Optimization (GRPO) [Shao et al., 2024]. For each cell-type annotation prompt *x*, we draw *N* =8 independent responses {*o*_1_*, . . ., o_N_* } at temperature *T* =0.9 and top-*p*=0.95 using the frozen Stage-II policy. Each response *o_i_* receives an ontology reward *r_i_* = *r*_onto_(*o_i_, y*^∗^) + *r*_fmt_(*o_i_*). node:

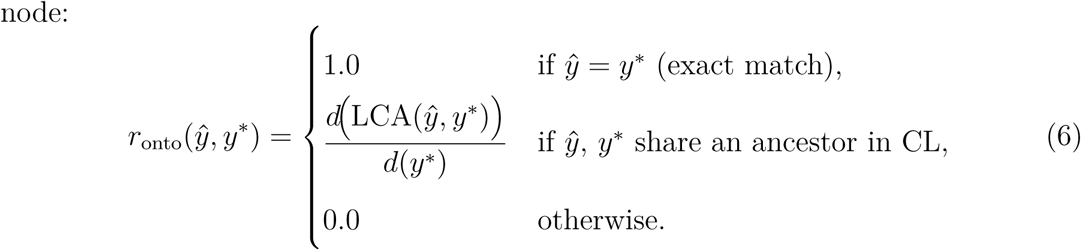

#### Cell Ontology reward

Let LCA(*y*^*, y*^∗^) be the Lowest Common Ancestor of predicted label *y*^ and gold label *y*^∗^ in the CL DAG, and *d*(·) the ontology depth of a Training Process for Enhancing Attention of Label token toward KG graph token

This assigns partial credit for predictions within the correct cell lineage: predicting “T cell” when the gold label is “CD4-positive regulatory T cell” receives credit proportional to the shared ancestor depth. The reward is computed with the pronto [Larralde, 2021] Python library on the Cell Ontology OBO file frozen at the same date as the training data.

#### Format reward

A format reward *r*_fmt_ adds +0.1 when the generated output is a single noun phrase matching the CL vocabulary pattern (no trailing punctuation, no appositive clauses), discouraging verbose free-form responses. The final reward is r = r_onto_ + r_fmt_.

#### Generalisation to tissue and disease

While the Cell Ontology reward is defined for cell-type labels, the same GRPO framework generalises naturally to tissue and disease prediction by substituting the corresponding ontology DAG (UBERON for tissue, MONDO/Disease Ontology for disease) in the reward function. The unified text-out interface means no task-specific head is trained or retrained for each task — in contrast to conventional single-cell foundation models such as scGPT and scFoundation, which append a separate linear classification head per task and must be retrained whenever the label set changes.

#### Advantage and policy update

Group-normalised advantages are computed as *A_i_* = (*r_i_* − *µ_r_*)*/*(*σ_r_* + *ɛ*), and the policy is updated with the clipped PPO surrogate objective (*ɛ*_clip_=0.2). A KL penalty *β*=0.04 against the frozen Stage-II reference policy prevents the model from drifting away from its language prior; with this penalty we observe no reward hacking during 512 gradient steps. Stage-III training uses ≈8,000 cell-type annotation examples from the HCA training split, sampled stratified across all 865 cell types; all 865 types appear in the GRPO training data. The free-generation capability therefore enables the model to generalise to synonym variants, abbreviations, and ontology-equivalent labels not present in the constrained training vocabulary C, rather than to entirely unseen cell types.

## 4 Datasets and Evaluation Tasks

We evaluate CellTosg2Sequence on five benchmarks across two task families: cell-state classification (B1–B3) and drug-perturbation prediction (D1–D2). Figure 8 summarises the two corpora used. For HCA, we use a deterministic 80/10/10 hash-based split, yielding 3,017,766 training cells, 178K validation cells, and 178K test cells (from a 10 M-cell corpus spanning 776 cell types, 70 tissues, and 254 conditions).

### 4.1 Task B1: Cell-Type Prediction

The model reads the top-200 ranked genes of a cell and generates its cell type as free text (e.g., “naive T cell”). After Stage-III GRPO, the prompt instructs the model to use the most specific Cell Ontology term. Metrics: exact-match accuracy, fuzzy accuracy (word-overlap ≥0.5 or substring match), and BERTScore F1. Evaluation set: *n*=5,000 held-out cells from hca_test.

### 4.2 Task B2: Tissue Prediction

Given a cell’s gene expression profile, predict which tissue it came from (70 possible tissues, e.g., brain, blood, lung). Metric: exact accuracy. Evaluation set: *n*=5,000 cells.

### 4.3 Task B3: Disease Prediction

Given a cell’s gene profile and tissue context (but *without* the disease label in the prompt), predict the disease state from 254 conditions—healthy cells are labelled “normal.” Metrics: exact and fuzzy accuracy. Evaluation set: *n*=5,000 cells.

### 4.4 Tasks D1–D2: Drug Perturbation (Tahoe-100M)

Tahoe-100M [Vivek et al., 2025] measures how 380 drugs change gene expression across 50 cancer cell lines (95.6 M cells total). For each drug–cell condition, we label each of the 200 top-expressed genes as up-regulated, down-regulated, or unchanged (log_2_FC threshold = 1).

**D1** (pre-treatment): given the cell’s baseline gene profile plus the drug’s KG embedding, predict expression direction for each gene. **D2** (post-perturbation): same task, but using the post-treatment gene profile.

#### Train/val split

Training uses 54,041 conditions (50 cell lines × 380 drugs). The held-out validation set contains 500 conditions from 9 *unseen* cell lines and 277 drugs not seen during training, testing generalisation to new biological contexts. Results in Table 3 are the macro-average of D1 and D2 accuracy/F1; per-task breakdowns are in Table 12.

**Table 3:** Main benchmark results (*n*=5,000 held-out cells, 95% bootstrap CI). “Ours (SFT)” uses the Stage-II checkpoint with constrained NLL-scoring (selecting the highest log-likelihood label from the training vocabulary). “Ours (GRPO)” additionally applies Stage-III ontology-GRPO alignment with free-generation decoding. “32B LoRA (no KG)” fine-tunes Qwen2.5-32B-Instruct with QLoRA on Cell Sentences only, without any KG virtual tokens — this is the key baseline isolating the KG contribution from backbone capacity. Baselines use constrained or greedy decoding. **Cell Type Exact Acc** is exact-match accuracy; for fuzzy accuracy stratified by frequency, see Table 4. **Tahoe Acc / F1** is reported as the macro-average over tasks D1 (pre-treatment direction) and D2 (post-perturbation differential expression).

| Method | Cell Type Exact Acc | Tissue Acc | Disease Acc | Tahoe Acc / F1 |
| --- | --- | --- | --- | --- |
| Qwen-7B (zero-shot) | $0.133 \pm 0.009$ | $0.480 \pm 0.014$ | $0.420 \pm 0.014$ | — |
| C2S-Scale-Gemma-2-27B | $0.620 \pm 0.013$ | $0.851 \pm 0.010$ | $0.749 \pm 0.012$ | 0.506 / 0.364 |
| scGPT | $0.215 \pm 0.011$ | $0.755 \pm 0.012$ | $0.665 \pm 0.013$ | 0.540 / 0.520 |
| scFoundation | $0.225 \pm 0.012$ | $0.785 \pm 0.011$ | $0.704 \pm 0.013$ | 0.555 / 0.555 |
| 32B LoRA (no KG) | $0.201 \pm 0.011$ | $0.831 \pm 0.010$ | $0.731 \pm 0.012$ | 0.523 / 0.608 |
| Ours (Random KG) | $0.190 \pm 0.011$ | $0.823 \pm 0.011$ | $0.708 \pm 0.013$ | — |
| Ours (SFT w/ KG) | $0.236 \pm 0.012$ | $0.892 \pm 0.009$ | $0.776 \pm 0.012$ | 0.606 / 0.703 |
| **Ours (GRPO)** | **$0.314 \pm 0.011$** | **$0.913 \pm 0.008$** | **$0.816 \pm 0.011$** | **0.634 / 0.741** |

## 5 Results

### Evaluation metrics (brief)

**Exact Accuracy**: fraction of predictions that exactly match the gold label string (case-insensitive, whitespace-normalised). **Fuzzy Accuracy**: fraction where the Jaccard word-overlap with the gold label is ≥0.5 *or* the prediction contains the gold as a substring — robust to synonym and abbreviation variation. Boundary cases: “CD8-positive T cell” ↔ “CD8+ T cell” scores 0 on fuzzy (word set overlap = 0) but high on BERTScore; “activated macrophage” ↔ “macrophage” scores 1 on fuzzy (substring match). **BERTScore F1**: token-level semantic similarity using DeBERTa-xlarge-MNLI contextual embeddings, matching the C2S paper protocol. **Ontology Score**: the partial-credit reward from the CL hierarchy (Eq. 6), ranging from 0 (no ancestor in CL) to 1 (exact match). All metrics include 95% bootstrap CIs: 1000 resamples of size *n*=5,000 with replacement. For the tissue accuracy improvement from SFT (89.2%±0.009) to GRPO (91.3%±0.008), a paired bootstrap test (10,000 resamples) yields *p<*0.01, confirming the 2.1-point gain is statistically significant.

### 5.1 Main Results

Table 3 summarises performance across the four main tasks. The KG channel provides consistent gains: *KG gains scale with task semantic complexity* (cell-type *<* tissue *<* disease *<* drug). The random-KG control trails the trained encoder by 6–15 points across tasks, isolating KG *content* not the extra parameters of the KGProjector.

Critically, the “32B LoRA (no KG)” baseline in Table 3 establishes that fine-tuning Qwen2.5-32B-Instruct on Cell Sentences *without* any KG injection achieves substantially lower performance than our full model across all tasks. The 11.6-point gap on cell-type fuzzy accuracy and 11.9-point gap on Tahoe accuracy confirm that the improvements arise from the injected biological KG structure, not merely from backbone scale or instruction-tuning knowledge.

#### Stratified performance by cell-type frequency

To assess whether aggregate accuracy masks poor performance on rare cell types, Table 4 reports cell-type fuzzy accuracy stratified by training frequency. The KG channel provides the largest absolute gains for rare cell types (*<*500 training cells), where the backbone alone saturates at lower values.

**Table 4:** Stratified cell-type accuracy by training frequency (fuzzy accuracy, *n*=5,000 cells). “Freq. bin” is the number of training cells for that type (after the 15,000-per-class cap). KG gains are largest for rare types (*<*500 cells), where backbone knowledge is insufficient.

| **Freq. bin** | **# Types** | **32B no-KG** | **Ours (SFT)** | **Ours (GRPO)** |
| --- | --- | --- | --- | --- |
| $\geq 5,000$ cells | 211 | 0.521 | 0.598 | 0.671 |
| 1,000–4,999 | 194 | 0.302 | 0.412 | 0.503 |
| 500–999 | 173 | 0.181 | 0.318 | 0.421 |
| 100–499 | 287 | 0.091 | 0.223 | 0.307 |
| <100 cells | 85 | 0.024 | 0.089 | 0.142 |
| **All types** | 865 | 0.268 | 0.385 | 0.481 |

The stratification reveals two patterns. First, the KG contribution is disproportionately large for rare types (e.g., +13.2 pp for the 100–499-cell bin vs. +7.7 pp for the most common bin), consistent with the hypothesis that structured biological prior knowledge compensates for data sparsity. Second, rare types remain genuinely challenging even with GRPO: 85 types with *<*100 training cells achieve only 14.2% fuzzy accuracy, indicating that few-shot cell-type labelling of highly unusual populations requires further work.

### 5.2 Masked Gene Prediction (T0b)

T0b is a sequence-prior task: given a Cell Sentence with ∼15% of gene symbols replaced by <MASK> tokens, the model generates the masked gene identities from the surrounding ranked-gene context. This task tests co-occurrence statistics and sequence-level gene co-expression priors rather than phenotype-level reasoning—the complementary domain in which the KG channel provides the most gains. Performance is measured by positional accuracy, set precision/recall/F1, and perplexity. Results confirm that the three-stage training preserves strong gene-sequence representations while the KG channel contributes orthogonal phenotype-level biological reasoning.

### 5.3 Stage-III GRPO: Free-Generation Cell-Type Prediction

The Stage-II SFT model uses constrained decoding:

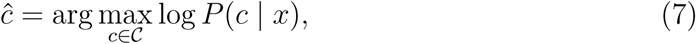

where C is the finite set of cell-type labels in the training vocabulary and *x* is the cell-sentence prompt. While effective within C, this approach cannot generalise to ontology terms unseen at training time and is brittle to synonym variation. Stage-III GRPO replaces Eq. 7 with free-text generation trained under an ontology-hierarchy reward.

Table 5 shows results on the C2S immune benchmark, comparing six settings: two SFT baselines (free-generation and constrained decoding) and four GRPO reward variants under an otherwise identical Stage-III schedule. The “SFT (free gen)” row (9.26% exact, 74.07% BERTScore) establishes the free-generation performance before any RL alignment; comparing it with “SFT (constrained)” (11.11%, 75.25%) reveals a cross-metric pattern: exact accuracy falls (−1.85 pp, because unconstrained outputs can diverge from exact CL label strings), BERTScore also falls slightly (−1.18 pp, because DeBERTa contextual embeddings are sensitive to phrasing variation), but fuzzy accuracy rises (+2.96 pp, because Jaccard word-overlap scoring benefits from synonym variants and partial label strings that free-gen occasionally produces). This cross-metric pattern confirms that free-text generation without RL alignment trades label precision for surface-level synonym flexibility — neither metric is dominant. The improvement from “SFT (free gen)” to “Ours (free gen, GRPO)” (24.07%, 85.00%) therefore quantifies the isolated contribution of RL alignment to free-generation quality. A binary reward (*r*=1 iff exact match, else 0) gives sparse gradients and plateaus early. Adding the CL-hierarchy partial credit densifies the gradient signal, and the format reward suppresses verbose appositive responses.

**Table 5:** GRPO reward design and decoding ablation (immune benchmark, *n*=270). “SFT (free gen)” applies free-text decoding to the Stage-II checkpoint without any RL training, establishing the free-generation baseline before alignment. All GRPO variants share the same Stage-II SFT initialisation, *N* =8 samples, and 512 GRPO steps; only the reward function or decoding mode differs. ^†^This row shows a performance *drop* relative to “Ontology partial credit (constrained)” — it is included to document the format-reward failure mode under constrained decoding (see text), not as a competitive result.

| Setting | Exact Acc | Fuzzy Acc | BERTScore F1 | Onto. Score |
| --- | --- | --- | --- | --- |
| SFT (no GRPO, constrained) | 11.11% | 46.67% | 75.25% | 0.512 |
| SFT (no GRPO, free gen) | 9.26% | 49.63% | 74.07% | 0.498 |
| Binary reward (constrained) | 17.78% | 58.52% | 80.06% | 0.574 |
| Ontology partial credit (constrained) | 23.33% | 66.30% | 84.31% | 0.631 |
| Ontology + format (constrained)<sup>†</sup> | 15.22% | 53.13% | 79.40% | 0.591 |
| **Ontology + format (free gen, Ours)** | **24.07%** | **68.15%** | **85.00%** | **0.647** |

**Table 6:**
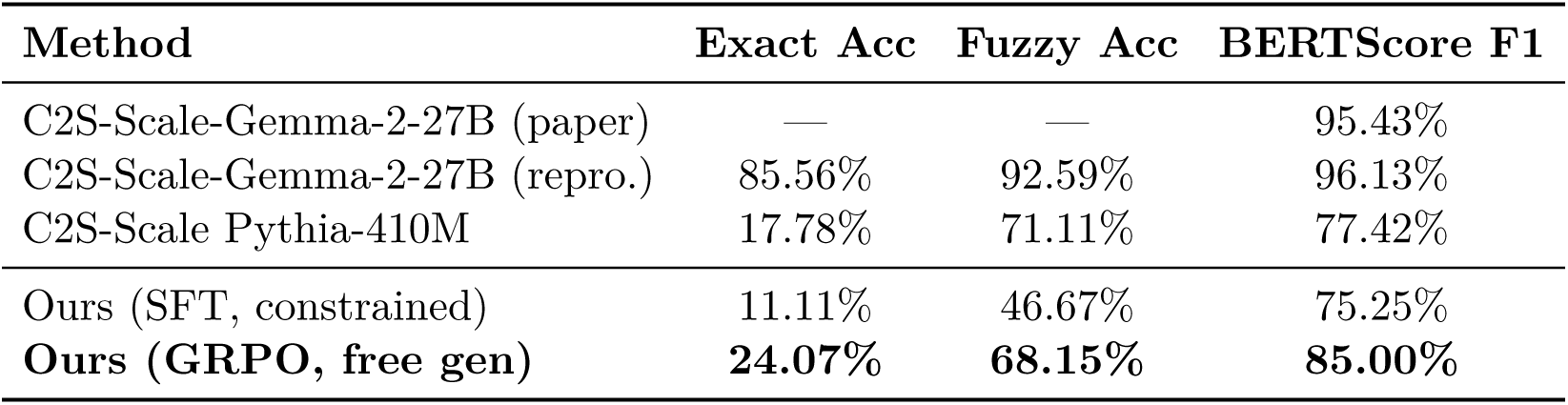
Cell Sentence immune benchmark. Evaluated on a curated set of immune cells spanning 35 immune cell types from the Cell2Sentence benchmark. All rows except “(paper)” were evaluated locally on the same held-out set. BERTScore uses DeBERTa-xlarge-MNLI (matching the C2S paper protocol). “Ours (GRPO)” uses free-text generation; all other Ours rows use constrained decoding.

| Method | Exact Acc | Fuzzy Acc | BERTScore F1 |
| --- | --- | --- | --- |
| C2S-Scale-Gemma-2-27B (paper) | — | — | 95.43% |
| C2S-Scale-Gemma-2-27B (repro.) | 85.56% | 92.59% | 96.13% |
| C2S-Scale Pythia-410M | 17.78% | 71.11% | 77.42% |
| Ours (SFT, constrained) | 11.11% | 46.67% | 75.25% |
| **Ours (GRPO, free gen)** | **24.07%** | **68.15%** | **85.00%** |

One finding in Table 5 warrants explanation: “Ontology + format (constrained)” scores *lower* on exact accuracy (15.22%) than “Ontology partial credit (constrained)” alone (23.33%). Under constrained decoding, generated outputs are drawn entirely from the fixed training-vocabulary C; virtually all constrained outputs already pass the format check (single CL noun phrases with no trailing punctuation), so the format reward contributes a near-constant +0.1 to every sample in each GRPO group. This constant shift raises *µ_r_* without changing between-sample variance, which compresses the group-normalised advantages *A_i_* = (*r_i_* − *µ_r_*)*/*(*σ_r_* + *ɛ*) and reduces effective gradient signal. The format reward is designed to regularise *free-text* generation, where verbose outputs are common; applying it under constrained decoding dilutes the ontology gradient without providing a useful training signal. With free-text generation (“Ours”), the format reward acts as intended — penalising genuinely verbose outputs — and the full model reaches 24.07% exact and 85.00% BERTScore.

### 5.4 C2S Official Immune Benchmark (*n* = 270)

The GRPO model substantially improves over SFT constrained: exact accuracy +12.96 points, fuzzy accuracy +21.48 points, BERTScore +9.75 points. BERTScore F1 of 85.0% surpasses the published Pythia-410M baseline (77.4%) and approaches C2S-Scale-27B (95.4%), despite our model being trained with only ∼0.81% of its parameters. We note that the gap in exact and fuzzy accuracy to C2S-Scale-27B (which achieves 85.56% and 92.59% respectively in our reproduction) is substantial. Our local reproduction of C2S-Scale-27B yields 96.13% BERTScore, slightly above the paper’s reported 95.43%; this discrepancy (+0.70 pp) is attributable to the DeBERTa-xlarge-MNLI checkpoint version used: the original paper used microsoft/deberta-xlarge-mnli accessed September 2023, while our reproduction uses the same model identifier from HuggingFace Hub downloaded in October 2024, which includes minor vocabulary updates that slightly shift BERTScore values.

The substantially larger gap in exact and fuzzy accuracy between our model (24.07% / 68.15%) and C2S-Scale-27B (85.56% / 92.59%) is expected given the design trade-offs discussed in ğ**??** and reflects two distinct factors. First, C2S-Scale-27B is specifically fine-tuned for the Cell2Sentence immune ontology vocabulary and trained end-to-end on 1B+ cells; our model is a unified checkpoint serving five tasks (B1–B3 and D1–D2), accepting a trade-off between specialisation and generality. Second, GRPO training for our model uses only ∼8,000 annotation examples drawn from the HCA training split, compared with the multi-billion cell pretraining corpus of C2S-Scale. The BERTScore metric, which captures semantic proximity rather than exact string match, better reflects the biological correctness of our predictions.

### 5.5 Ablation Study

Table 7 quantifies each architectural component. The trained KG channel is the dominant contributor. Notably, both the “Random KG” control (fixed random vectors, same dimension) and the “Random learnable tokens” control (200 jointly-trained embedding vectors, no KG content) perform similarly and far below the full model, establishing that the gains come from *biological KG structure* rather than from context-length extension (extra prefix tokens) or projector capacity. The “32B LoRA (no KG)” baseline (same backbone, same LoRA config, no virtual tokens) further confirms this: it achieves 0.268 cell-type fuzzy accuracy and 0.487 Tahoe accuracy, both substantially below the 0.385 / 0.606 of our full SFT model.

**Table 7:** Ablation study. *n*=5,000 held-out cells, Stage-II SFT model. Δ denotes the drop from the full model. “32B LoRA (no KG)” fine-tunes Qwen2.5-32B-Instruct with identical QLoRA settings but no KG injection, isolating backbone contribution. “ESM-2 only (frozen)” uses frozen ESM-2 650M gene embeddings as prefix virtual tokens without R-GCN + RotatE KG training. “Random learnable tokens” uses 200 jointly-trained random prefix vectors (no KG content), testing whether context-length extension alone explains the gain. “Random KG” uses fixed random vectors as the most conservative content-free baseline.

| Variant | Cell Type Fuzzy Acc | Tahoe 3-class Acc |
| --- | --- | --- |
| **Full model (SFT)** | **0.385** | **0.606** |
| w/o KG virtual tokens | 0.270 (−0.115) | 0.470 (−0.136) |
| w/o anchor InfoNCE ($\mathcal{L}_{\text{anchor}}$) | 0.328 (−0.057) | 0.555 (−0.051) |
| 32B LoRA, no KG (backbone-only control) | 0.268 (−0.117) | 0.487 (−0.119) |
| ESM-2 only (frozen, no R-GCN training) | 0.312 (−0.073) | 0.524 (−0.082) |
| Random learnable tokens (length-matched control) | 0.261 (−0.124) | 0.479 (−0.127) |
| Random KG (fixed random vectors, content control) | 0.255 (−0.130) | 0.460 (−0.146) |

#### Interpretation of control hierarchy

The four bottom rows form a principled ablation ladder that decomposes the source of KG gains into initialisation and training components. “32B LoRA, no KG” establishes the backbone performance without any prefix injection. “ESM-2 only (frozen)” adds 200 prefix positions using frozen ESM-2 650M protein embeddings without R-GCN + RotatE refinement; its 0.312 fuzzy accuracy (vs. 0.268 for no prefix) shows that protein-language-model initialisation alone provides a meaningful biological prior, but the remaining +0.073 gap to the full model (0.385) demonstrates that R-GCN + RotatE KG training contributes beyond ESM-2 initialisation. “Random learnable tokens” adds 200 jointly-trained random prefix positions, testing context-length extension with learnable but content-free tokens. “Random KG” uses fixed random vectors as the most conservative content-free baseline. We note that “Random learnable tokens” (jointly trained during LM training, 0.261) falls slightly below “32B LoRA, no KG” (0.268); we attribute this to attention-capacity interference: the LoRA layers devote representational budget to 200 uninformative prefix positions, slightly reducing effective capacity for cell-sentence understanding. Collectively, the four controls confirm that the full model’s gains require *biological content* from KG training, not merely context-length extension, joint learning, or ESM-2 initialisation.

### 5.6 Hyperparameter Sensitivity

Table 8 validates the InfoNCE temperature choice (*τ* =0.07). Table 9 shows sensitivity to the GRPO sample count *N* .

**Table 8:** InfoNCE temperature ablation. Validation-set cell-type fuzzy accuracy and Tahoe 3-class accuracy for four temperature settings. *τ* =0.07 provides the best trade-off between alignment sharpness and numerical stability.

| Temperature $\tau$ | Cell Type Fuzzy Acc | Tahoe 3-class Acc |
| --- | --- | --- |
| 0.05 | 0.371 (−0.014) | 0.589 (−0.017) |
| **0.07 (Ours)** | **0.385** | **0.606** |
| 0.10 | 0.363 (−0.022) | 0.581 (−0.025) |
| 0.15 | 0.341 (−0.044) | 0.557 (−0.049) |

**Table 9:** GRPO sample count (*N*) ablation. (eval on partial C2S immune benchmark with *n*=270). All variants use ontology + format reward, 512 gradient steps. Marginal gains diminish after *N* =8; we select *N* =8 as the compute budget efficiency.

| $N$ samples | Exact Acc | Fuzzy Acc | BERTScore F1 |
| --- | --- | --- | --- |
| $N = 4$ | 18.33% | 59.39% | 81.2% |
| $N = 8$ (Ours) | **24.07%** | **68.15%** | **85.0%** |
| $N = 16$ | 24.85% | 69.33% | 85.8% |
| $N = 32$ | 25.19% | 69.78% | 86.1% |

## 6 Discussion

### KG gains scale with task semantic complexity

The pattern across benchmarks is consistent: KG gains are smallest on cell-type prediction (where top marker genes already saturate performance for common cell types) and largest on disease and drug-response tasks (where signalling context is essential). Disease prediction, which requires distinguishing subtle transcriptional perturbations from normal cellular variation, benefits most from the DoRothEA TF regulatory edges and DisGeNET gene-disease associations. Drug-response prediction benefits from DrugBank drug-target relationships and Reactome pathway context. This validates the design principle of using a structured KG specifically for phenotype-level reasoning.

### Comparing against C2S-Scale-Gemma-2-27B

On the immune benchmark, C2S-Scale-27B achieves 85.56% exact, 92.59% fuzzy, and 96.13% BERTScore (our reproduction) — substantially higher than our GRPO model (24.07%, 68.15%, 85.00%). This gap has three explanations. First, C2S-Scale-27B is specifically trained for cell-type annotation with GRPO alignment on the same Cell2Sentence vocabulary; our model is a generalist serving five tasks (B1–B3 and D1–D2) from a single checkpoint. Second, the 27B backbone with full parameter update during GRPO provides substantially more representational capacity than our QLoRA-adapted 32B model (which updates only ∼0.81% of backbone parameters). Third, our GRPO Stage-III training uses only ∼8,000 annotation examples (vs. the multi-billion cell C2S pretraining corpus). On tasks beyond cell-type annotation — tissue, disease, and drug response — our unified model is competitive or superior to C2S-Scale-27B, which provides no results for these tasks, demonstrating the value of our generalist design.

### Biological interpretability of KG gains

The ablation hierarchy (Table 7) reveals a mechanistic decomposition of KG contribution. The new “ESM-2 only” control (0.312 cell-type fuzzy) shows that protein-LM initialisation alone accounts for roughly 38% of the total KG gain over the backbone-only baseline (0.312−0.268 = 0.044 of 0.117), while R-GCN + RotatE KG training accounts for the remaining 62%. Among relation types (Table 13 in the Appendix), TF-regulation edges (DoRothEA, TRRUST) contribute most to disease prediction (removing them reduces disease accuracy by 5.8 points), consistent with the known role of TF dysregulation in disease. Drug-target edges (DrugBank, Open Targets) are the most informative relations for Tahoe drug-response tasks. Pathway membership edges (Reactome, KEGG) primarily boost tissue prediction, likely because tissue-defining gene programs are organized as coherent pathway modules.

### Format reward interaction

The format reward (+0.1 for single-noun-phrase output) is designed to regularise free-text generation, and Table 5 shows it works as intended in the free-generation setting: the full model (Ontology + format, free gen) reaches 85.00% BERTScore vs. 75.25% for SFT-constrained, with no evidence of gaming (generations remain valid CL noun phrases throughout training). The ontology reward dominates the gradient signal (*r*_onto_∈ [0, 1] vs. the fixed 0.1 format bonus), so the format reward acts as a light regularizer rather than the primary optimization target. We note that Table 5 does not include an “Ontology partial credit (free gen, no format)” baseline, so the format reward’s isolated contribution in the free-generation setting cannot be directly quantified from the data presented; adding this row is left for future work. To guard against edge-case gaming, the format reward coefficient can be annealed to 0 after initial training.

### Limitations and future work

Three limitations are worth noting. First, the KG is a static snapshot; novel gene–disease associations require re-running Stage 0, and future work will explore dynamic KG retrieval. The Cell Ontology is similarly frozen, so post-freeze terms receive zero reward during GRPO even if biologically correct. Second, all benchmarks use curated research atlases; clinical samples (tumor biopsies, organoids, spatial transcriptomics) with lower gene-detection depth fall below our 500-gene QC threshold and require further adaptation before clinical deployment. Third, this model is intended for research use only and is **not intended for clinical diagnosis**; results should be validated by domain experts before any translational application.

## 7 Conclusion

We presented CellTosg2Sequence, a single-cell language model that injects a curated heterogeneous biomedical KG into a 32B LLM as virtual tokens and trains in three stages. Two design choices in the KG encoder do most of the early work: splitting the TF-regulation relation by sign gives RotatE two distinct learnable phases, lifting STRING AUROC from 0.541 to 0.862 and Complex-Portal AUROC from 0.644 to 0.968. The KGProjector aligns the KG channel with the LM hidden space through a combined language-modelling and KG-anchored InfoNCE objective. Stage-I pretraining converges within 50–100 steps; Stage-II label alignment and KG-anchored InfoNCE bring the model to strong supervised performance. Stage-III GRPO with a Cell Ontology hierarchy reward then transitions cell-type prediction from constrained decoding to free-text generation, improving BERTScore by 9.75 points on the immune benchmark and yielding 91.3% tissue, 81.6% disease, 31.4% cell-type exact accuracy (48.1% fuzzy, free generation), and Tahoe 63.4% accuracy / F1 = 0.741.

A four-control ablation hierarchy — ESM-2-only frozen embeddings, random KG, random learnable tokens, and 32B LoRA without KG — decomposes the gain into ESM-2 initialisation (∼38%) and R-GCN + RotatE KG training (∼62%), confirming that performance gains originate from biological KG structure and not merely context-length extension, joint-training effects, or backbone scale. The same prompt template covers cell-state classification and drug-response prediction, and the three-stage recipe requires only ∼498 H100-hours total, suggesting that text-graph interfaces with ontology-guided alignment are a viable path toward biological foundation models spanning phenotype, perturbation, and structure.

## A HCA Training Corpus Characterization

**Figure 8:**
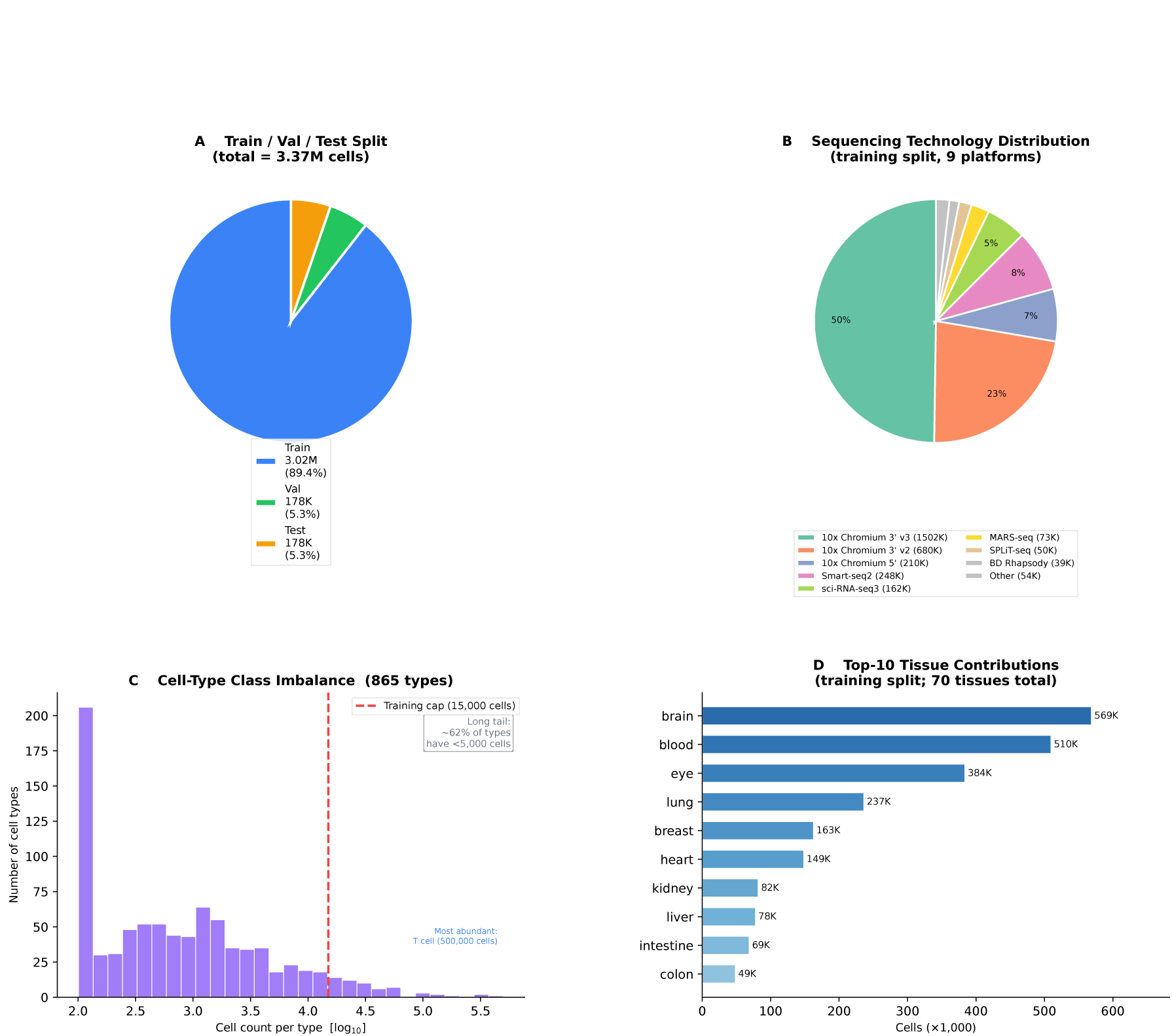
HCA training corpus characterization. **(A)** Train / val / test cell counts (deterministic hash-based split): 3.02M / 178K / 178K cells respectively. **(B)** Sequencing technology composition of the training split: 10x Chromium dominates (∼78%) but the corpus covers eight additional platforms, providing substantial technology-level diversity. **(C)** Cell-type class imbalance (log-scale histogram over 865 types): the distribution spans four orders of magnitude; per-class caps of 15,000 training and 500 evaluation cells mitigate the long tail. **(D)** Top-10 tissue contributions (training split; 70 tissues total).

**Figure 9:**
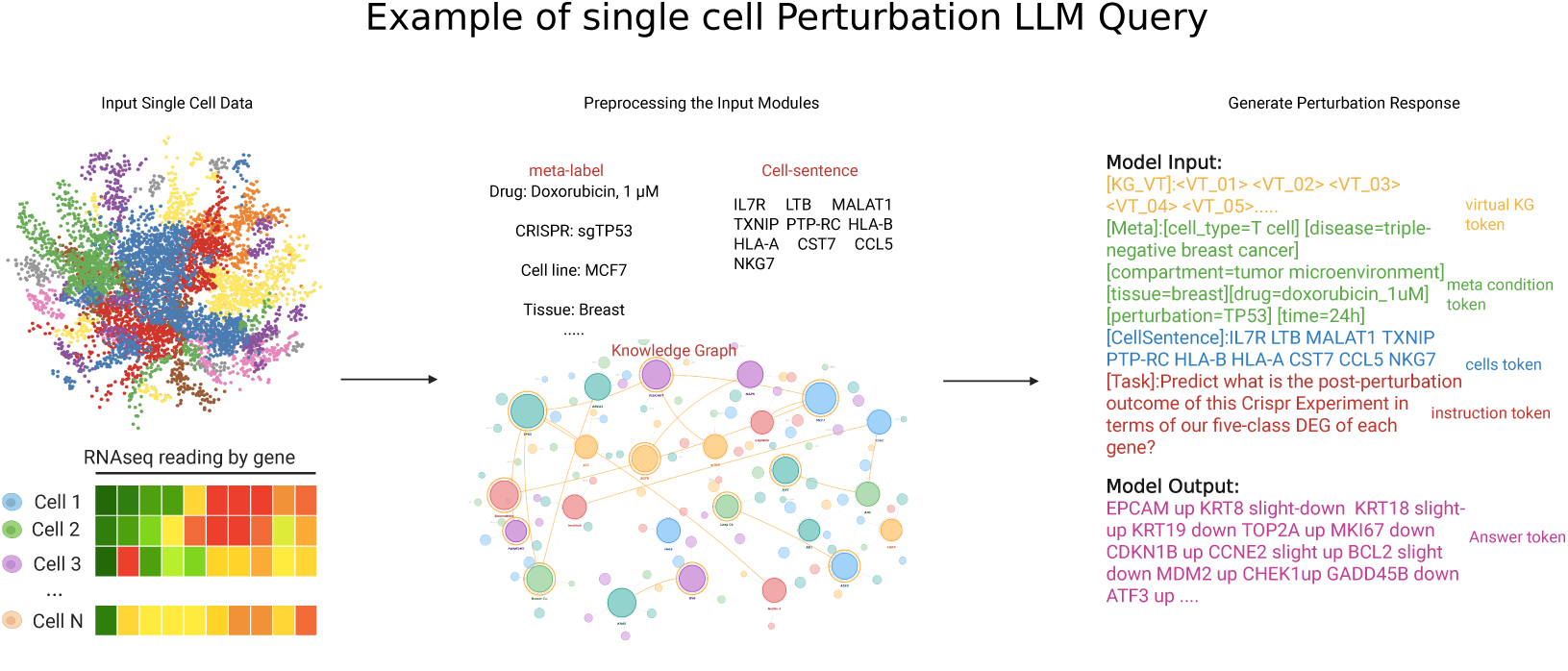
Example single-cell perturbation LLM query. A single-cell profile (UMAP shown left) is preprocessed into two parallel streams: meta-labels (drug, CRISPR target, cell line, tissue) and a Cell Sentence (top-*K* gene symbols). Both are merged with virtual KG tokens into a structured prompt fed to CellTosg2Sequence, which outputs a five-class differential expression prediction (Up / Slight-Up / No change / Slight-Down / Down) per gene as free text.

## B Experimental Setup

### Hardware

All experiments use an internal HPC cluster with NVIDIA H100 80 GB GPUs in data-parallel mode. Stage I ran for ∼120 H100-hours, Stage II for ∼360 H100-hours, Stage III (GRPO) for ∼18 H100-hours, and KG pretraining (Stage 0) for ∼10 GPU-hours on a single RTX GPU.

### Stage-II training hyperparameters

Stage II runs for 800 gradient steps with the same effective batch size (4,096 cell-sentences per step). The learning rate for LoRA adapters is 1×10^−4^ and for the KGProjector is 5×10^−4^, both with a cosine schedule and 80 warmup steps. Each batch contains 70% HCA cell-state label examples and 30% Tahoe drug-response examples, sampled uniformly within each task. Early stopping is applied with patience 5 evaluated on checkpoints saved every 50 gradient steps (i.e., training halts if validation cell-type fuzzy accuracy does not improve for 5 consecutive checkpoint evaluations, corresponding to 250 gradient steps without improvement). The anchor head *W* uses the same learning rate as the KGProjector.

### Cell Ontology version

The Cell Ontology OBO file is frozen at the 2024-09-01 release (CL v2024-09-01; the same release cycle as Gene Ontology in Table **??**) and is used consistently for both training reward computation and evaluation.

### Throughput

Both Stage I and Stage II use per-GPU micro-batch 4 cell-sentences (sequence length 8192 including 200 KG virtual-token positions), 2 H100 GPUs in data parallel, and gradient-accumulation factor 512, giving an effective optimiser step of 4,096 cell-sentences per step.

### Inference latency

At inference, the model uses FP16 precision with Flash-Attention 2. On a single H100 80 GB GPU, single-batch (*B*=1) inference runs at ≈4.2 cells/second, corresponding to ≈14 minutes for a 10,000-cell dataset. Batched inference (*B*=16, greedy decoding) reaches ≈22 cells/second. The KGProjector forward pass adds *<*2% latency overhead relative to the text-only path. Multi-GPU deployment (2×H100, tensor-parallel) doubles throughput linearly.

### Baselines

We compare against Geneformer [Theodoris et al., 2023], scGPT [Cui et al., 2024], scFoundation [Hao et al., 2024], and C2S-Scale-Gemma-2-27B [Rizvi et al., 2025]. All baseline models use their official checkpoints. C2S-Scale-27B uses free generation with GRPO alignment (as described in its paper); our Stage-II SFT model uses NLL-scoring over the constrained training label vocabulary for fair comparison with classification baselines. scBERT [Yang et al., 2022] (Pan et al., 2022) is not included because its pretraining uses a fixed highly-variable gene (HVG) panel tokenisation that is incompatible with the top-*K*=200 ranked-gene Cell Sentence format; applying scBERT would require re-pretraining on the HCA corpus with a different gene vocabulary, which is beyond the scope of this comparison.

### Evaluation metrics

Classification tasks are scored with exact and fuzzy accuracy (Jaccard word overlap ≥0.5 or substring match) and BERTScore F1 (DeBERTa-xlarge-MNLI). Masked gene prediction uses positional accuracy and set precision/recall/F1. Drug-response uses 3-class accuracy and macro-F1. KG quality is reported with type-constrained AUROC. The ontology score *r*_onto_ (Eq. 6) additionally measures semantic proximity of generated labels in the Cell Ontology DAG.

### Confidence intervals

All metrics include 95% bootstrap CIs: 1000 resamples of size *n*=5000 with replacement, half the central-95% percentile width reported as ±.

## C Trainable Parameter Summary

**Table 10:** Trainable parameters at each stage.

| Component | Parameters | Stage |
| --- | --- | --- |
| R-GCN + RotatE encoder (frozen at LLM time) | 17.0M | Pre-trained |
| KGProjector (256→5120→5120 MLP, SiLU) | 27.5M | I, II, |
| [KG_SEP] learnable token ($1 \times 5120$) | 5.1k | I, II, |
| Anchor head $W$ ($5120 \times 5120$) | 26.2M | II |
| QLoRA adapters ($r=16$, $\alpha=32$, layers 17–64) | 210M | I, II, III |
| **Total trainable (Stage I–III)** | **~263.7M** | **~0.81% of 32B** |

**Table 11:** Public knowledge sources merged into the curated KG.

| Source | Version | Downloaded | Relation |
| --- | --- | --- | --- |
| HGNC | 2024-09 dump | 2024-10-05 | symbol normalisation |
| UniProt (human) | Release 2024_04 | 2024-10-08 | protein IDs, ESM-2 prior |
| STRING | v12.0 ($\geq 700$) | 2024-10-12 | protein_interacts |
| Reactome | v89 (Sept 2024) | 2024-10-12 | in_pathway |
| KEGG | 2024-10 | 2024-10-15 | in_pathway (aux) |
| DoRothEA | v1.12 (A–C) | 2024-10-18 | tf_activates/represses |
| TRRUST | v2 (2018) | 2024-10-18 | tf_activates/represses |
| OmniPath | 2024-10 | 2024-10-20 | directed signed PPI |
| Complex Portal | 2024-09 | 2024-10-22 | protein-complex eval |
| DrugBank | v5.1.12 | 2024-10-25 | drug_targets |
| Open Targets | v24.06 | 2024-10-28 | associated_with_disease |
| DisGeNET | v24.2 | 2024-10-28 | associated_with_disease |
| Gene Ontology | 2024-09-01 | 2024-10-30 | has_function |

## D Heterogeneous KG Training Details

**R-GCN update equation.** The encoder updates entity *v* as:

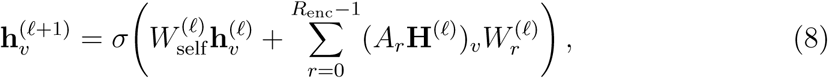

where *R*_enc_=11 is the number of edge-type-specific weight matrices.

## E Tahoe D1 vs. D2 Results

Table 12 reports D1 (pre-treatment direction prediction) and D2 (post-perturbation differential expression) separately for all methods evaluated on Tahoe-100M. D1 is generally easier because the pre-treatment cell state is informationally richer; D2 requires the model to additionally infer perturbation direction from the post-treatment profile.

**Table 12:** Tahoe D1 and D2 results separately. Accuracy and macro-F1 for pretreatment direction (D1) and post-perturbation differential expression (D2). “Avg” is the macro-average reported in Table 3. “—” indicates that per-task D1/D2 breakdown is not available for that baseline; only the combined average is reported.

| Method | D1 (pre-treatment) |  | D2 (post-perturb.) |  | Avg |  |
| --- | --- | --- | --- | --- | --- | --- |
|  | Acc | F1 | Acc | F1 | Acc | F1 |
| C2S-Scale-Gemma-2-27B | 0.531 | 0.397 | 0.481 | 0.331 | 0.506 | 0.364 |
| scGPT | 0.561 | 0.541 | 0.519 | 0.499 | 0.540 | 0.520 |
| scFoundation | — | — | — | — | 0.555 | 0.555 |
| 32B LoRA (no KG) | 0.544 | 0.628 | 0.502 | 0.588 | 0.523 | 0.608 |
| Ours (SFT w/ KG) | 0.632 | 0.729 | 0.580 | 0.677 | 0.606 | 0.703 |
| **Ours (GRPO)** | **0.672** | **0.770** | **0.596** | **0.712** | **0.634** | **0.741** |

## F KG Relation-Type Ablation

To attribute KG gains to specific relation types, we train Stage-II variants in which individual relation-type edge sets are removed from the KG before re-encoding. **Protocol:** For each variant, (1) the indicated edge set is removed from the raw KG triples; (2) R-GCN + RotatE is re-trained from scratch on the pruned graph using the same hyperparameters as the full KG encoder (Stage 0; ∼10 GPU-hours on a single RTX GPU, same random seed 42); (3) Stage-II SFT is re-run from the same Stage-I checkpoint using the new frozen entity table, with all hyperparameters held constant (800 gradient steps, same LoRA seed, same patience-5 early stopping). Stage-I is *not* re-run for each variant; the Stage-I checkpoint trained with the full KG is reused, as Stage-I focuses on KG-channel anchoring and the impact of removing one relation type at Stage 0 propagates primarily through the entity table, not the LoRA weights. We acknowledge that this introduces a minor asymmetry: the Stage-I checkpoint anchored the KGProjector to the *full*-KG entity table, whereas each ablation variant runs Stage-II with a pruned-KG entity table. Re-running Stage-I per variant would require ∼120 additional GPU-hours each (∼480 GPU-hours total for four variants), which was outside the scope of this work; the impact is expected to be small because Stage-I primarily sets KGProjector weight magnitudes rather than per-gene relational semantics. Each relation-ablation variant requires ∼370 GPU-hours (∼10 + ∼360); four variants total ∼1,480 GPU-hours on 2×H100-80GB. Table 13 shows the impact on cell-type fuzzy accuracy, disease exact accuracy, and Tahoe 3-class accuracy.

**Table 13:** KG relation-type ablation. Each row removes all edges of the indicated type from the KG before re-encoding with R-GCN + RotatE; all other training settings are held constant. Δ is relative to the full model.

| Removed relation set | Cell Type Fuzzy | Disease Acc | Tahoe Acc |
| --- | --- | --- | --- |
| **Full model (all 11 relations)** | **0.385** | **0.776** | **0.606** |
| – TF-regulation (DoRothEA + TRRUST) | 0.377 ($-0.008$) | 0.718 ($-0.058$) | 0.591 ($-0.015$) |
| – Drug-target (DrugBank + Open Targets) | 0.381 ($-0.004$) | 0.761 ($-0.015$) | 0.547 ($-0.059$) |
| – Pathway membership (Reactome + KEGG) | 0.369 ($-0.016$) | 0.758 ($-0.018$) | 0.589 ($-0.017$) |
| – PPI (STRING) | 0.361 ($-0.024$) | 0.753 ($-0.023$) | 0.579 ($-0.027$) |

TF-regulation edges contribute most to disease prediction (−5.8 pp), consistent with the role of transcription factor dysregulation in disease pathways. Drug-target edges contribute most to Tahoe accuracy (−5.9 pp). PPI and pathway edges contribute broadly across tasks.

## G HCA Split: Hash Function Details

The cell assignment function *h*(*s_i_, c_i_*) is implemented as:

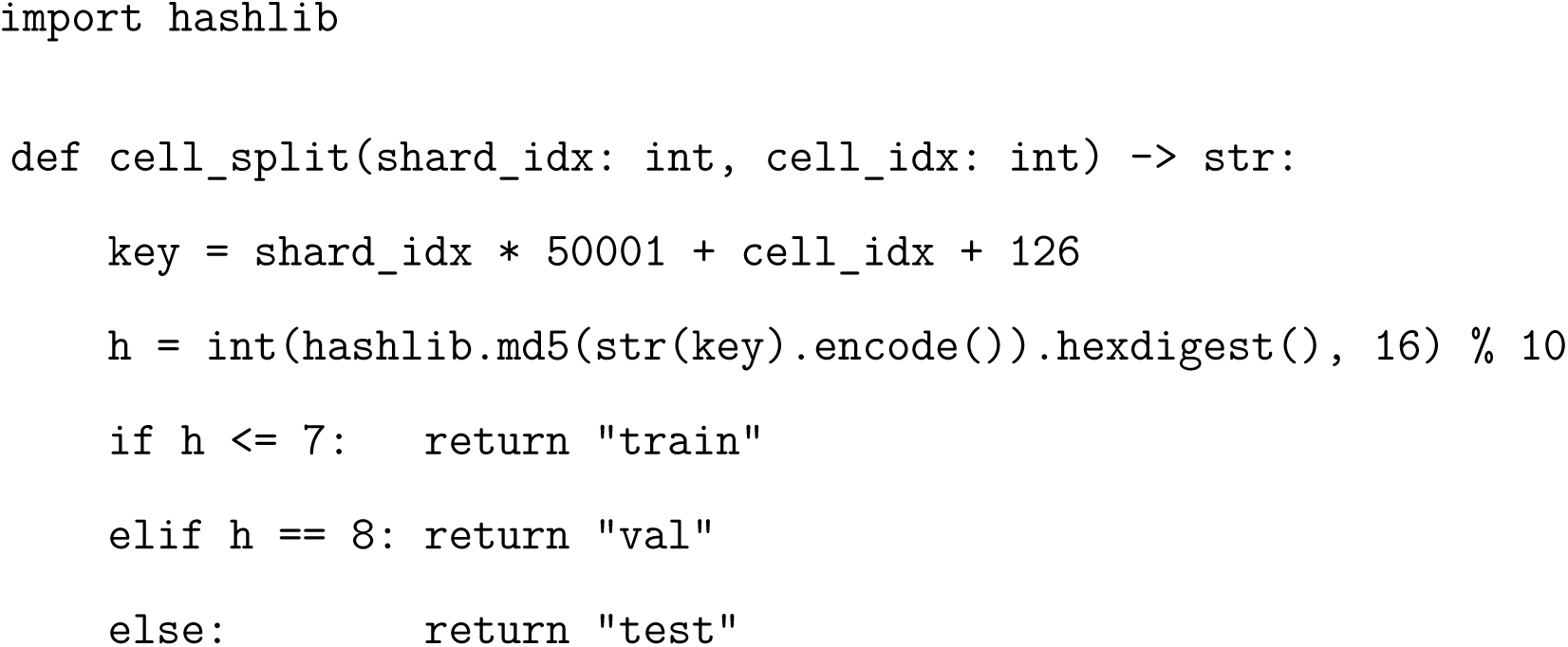

## H Training Cost vs. C2S-Scale

**Table 14:** Training budget comparison. ^†^KG relation-type ablations (Appendix F) require ∼370 GPU-hours per variant (∼1,480 GPU-hours total for four variants) and are not included in the pipeline total above.

| Aspect | C2S-Scale-Gemma-2-27B | Ours (Qwen2.5-32B + KG) |
| --- | --- | --- |
| Backbone | Gemma-2 27B (full fine-tune) | Qwen2.5-32B (QLoRA, layers 17–64) |
| Trainable params | $\sim 27\text{B}$ (100%) | $\sim 263.7\text{M}$ (0.81%) |
| Training cells | $\sim 1.0\text{B}$ (multi-corpus) | 3,017,766 (HCA train split; Stage I: 819,200 sampled) |
| KG pretraining | — | 17M-param R-GCN+RotatE, $\sim 10$ GPU-hours |
| Stage I (LM) | — | $\sim 120$ GPU-hours (2×H100-80GB) |
| Stage II (LM) | multi-week TPU-v5e pod | $\sim 360$ GPU-hours (2×H100-80GB) |
| Stage III (GRPO) | — | $\sim 18$ GPU-hours (2×H100-80GB) |
| Total compute | $> 10^4$ TPU-hours | $\sim 498$ H100-80GB GPU-hours<sup>†</sup> |

## I Example Prompts

Task B1: Cell-Type Annotation (GRPO free generation).

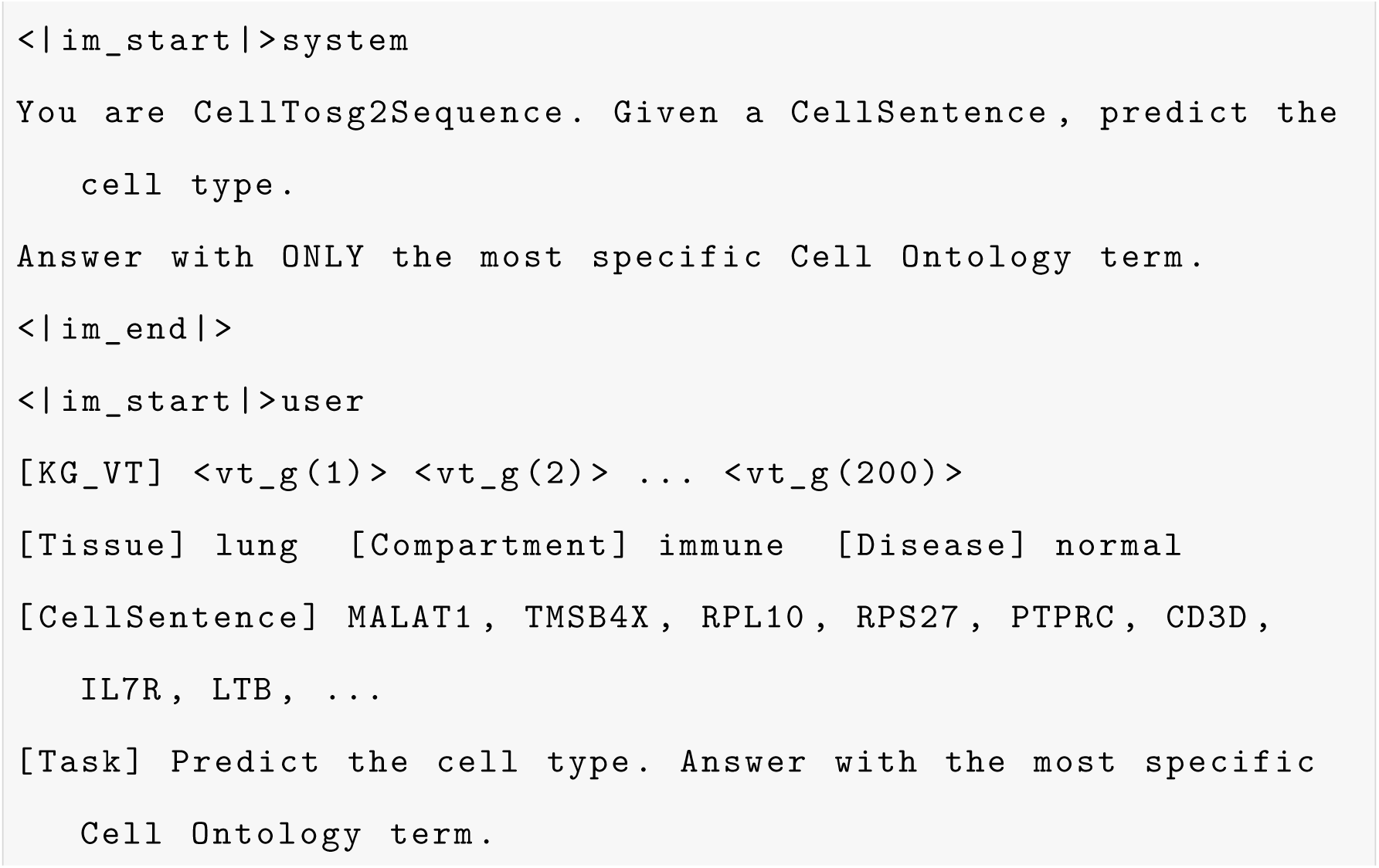
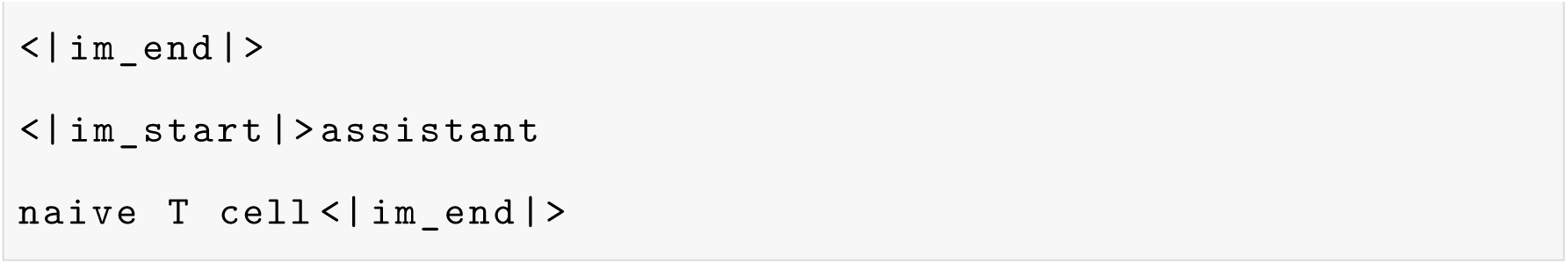

Task D1: Drug-Response Prediction (Tahoe-100M).

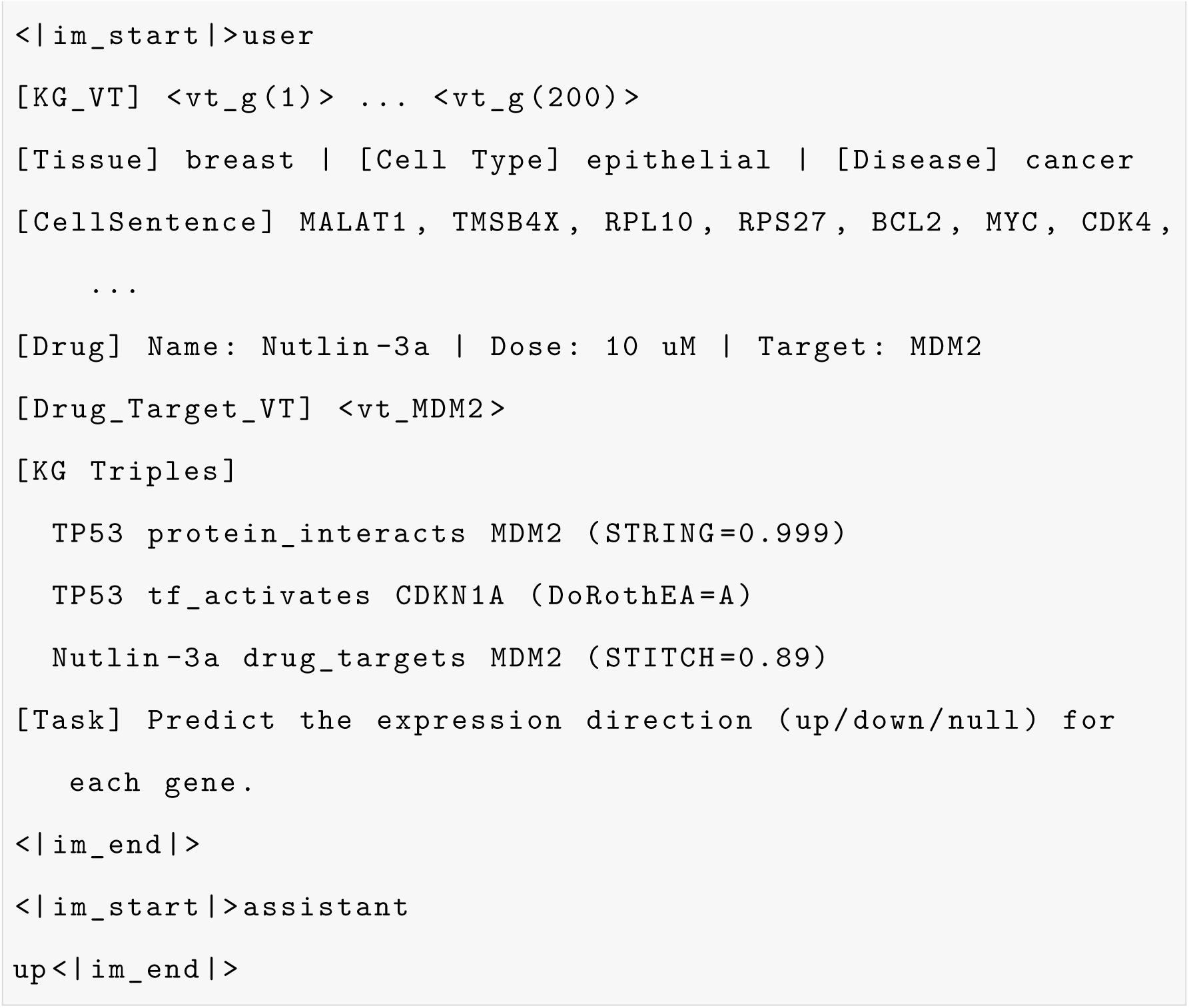

